# The Pregnane X Receptor (PXR) contributes to phenotypic plasticity and modulates the sensitivity to targeted therapies in melanoma

**DOI:** 10.64898/2026.11.03.762858

**Authors:** Stefan D. Nicolescu, Alice Matheux, Marina Grimaldi, Nadia Vie, Raphaël Romero, Mathilde Robin, Eric Fabbrizio, Abdelhay Boulahtouf, Julia Toucheboeuf, Amélie Gudin-De-Vallerin, Florence Boissière, Fanny Leenhardt, Litaty Mbatchi, Laura Tallevast, Patrick Balaguer, Céline Gongora, Alexandre Evrard, Philippe Pourquier

**Author notes:** Contributed equally (Alexandre Evrard, Philippe Pourquier).

## Abstract

The challenge in treating advanced metastatic melanoma lies in the development of therapeutic resistance and the risk of recurrence. Understanding the mechanisms underlying tumor adaptation to treatment is therefore crucial, as they can directly influence therapeutic efficacy. Tolerance to targeted therapies may be influenced by drug transport and metabolism within tumors, processes regulated by numerous genes under the transcriptional control of the nuclear receptor PXR (Pregnane X Receptor, NR1I2). We revealed, for the first time, significant and heterogeneous expression of PXR in human melanoma samples. We previously showed that PXR can be activated by BRAF inhibitors (BRAFi), but not by MEK inhibitors (MEKi), which are currently used in combination for the treatment of BRAF-V600E-mutated melanoma. Here, we investigated whether PXR expression influences melanoma cell sensitivity to MAPK pathway inhibitors. Using BRAF-mutated human melanoma cell lines, we showed that stable PXR expression sensitizes cells to combined BRAFi/MEKi treatment. Transcriptomic analysis further revealed that PXR expression is associated with the upregulation of several phenotypic markers, with MITF identified as a major regulator in untreated PXR-overexpressing cells. These findings suggest that PXR promotes a shift toward a melanocytic phenotype, which is associated with increased sensitivity to BRAFi/MEKi treatment. Characterization of the PXR-associated transcriptional signature in melanoma may therefore have translational value for identifying patients more likely to benefit from targeted therapies and support the development of PXR as a predictive biomarker of therapeutic response.

**GRAPHICAL ABSTRACT:** 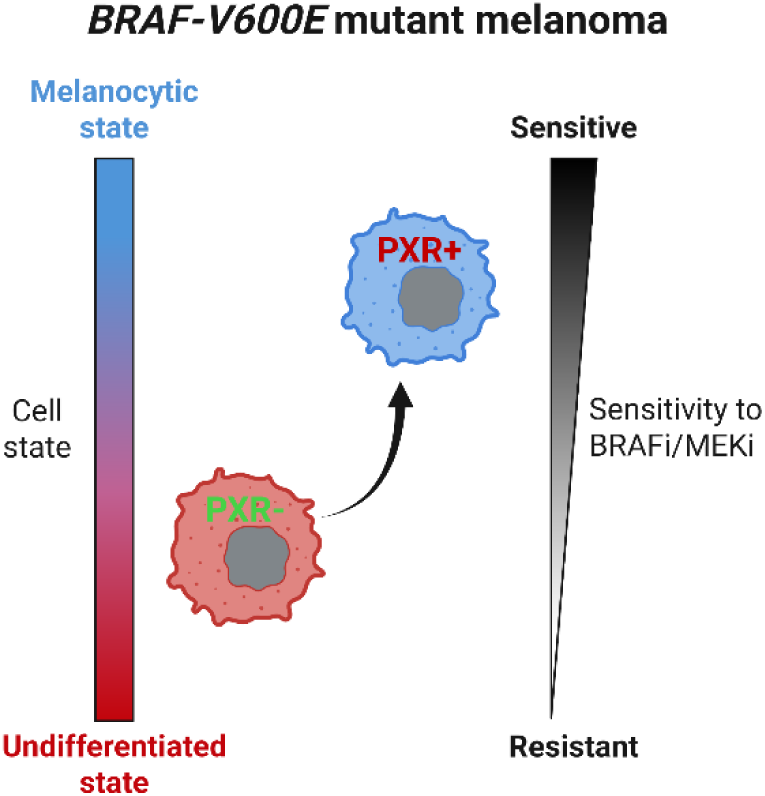

## INTRODUCTION

Cutaneous melanoma is a relatively uncommon skin cancer but is potentially aggressive and responsible for the majority of skin cancer-related deaths^1^. In 2024, an estimated 337,969 new melanoma cases were diagnosed and 58,398 deaths were reported worldwide^2^. Melanoma is characterized by marked cellular plasticity and a high mutational burden, contributing to its aggressiveness.

Major therapeutic advances, particularly the introduction of BRAF inhibitors (BRAFi: dabrafenib, encorafenib, or vemurafenib) combined with MEK inhibitors (MEKi: trametinib, binimetinib, or cobimetinib) for patients harboring a BRAF-V600 mutation (>50%), as well as immune checkpoint inhibitors, have substantially improved patient survival^3^. Nevertheless, their long-term efficacy is limited by the development of intrinsic or acquired resistance, and a considerable proportion of patients ultimately experience disease progression^4^. Understanding the molecular mechanisms involved in tumor cell response to these targeted therapies is therefore essential for improving treatment efficacy and for the identification of biomarkers that could help to select patients who would benefit most from these treatments.

The resistance to BRAF or MEK inhibitors is predominantly associated with the reactivation of the MAPK pathway that involves BRAF amplification or alternative splicing, or mutations in NRAS, BRAF, or MEK^3,5–7^. It may also involve activation of the alternative PI3K-AKT-mTOR pathway^8–10^. Non-genetic mechanisms involving epigenetic changes that promote tumor cell plasticity can emerge during tumor recurrence, enabling the reprogramming of oncogenic signaling pathways and cellular metabolism that ultimately reduce treatment efficacy^6,11–13^. Therapeutic response may also depend on drug metabolism or transport that is regulated by numerous genes under the transcriptional control of the nuclear receptor PXR (Pregnane X Receptor), encoded by the NR1I2 gene^14^. Upon activation by its ligands, PXR forms a heterodimer with RXR and binds to regulatory regions of target genes^15–17^. These include genes encoding efflux transporters such as P-Glycoprotein (ABCB1)^16,18,19^ or Breast Cancer Resistance Protein (ABCG2)^19^, as well as metabolic enzymes such as cytochrome P450 3A4 (CYP3A4)^16,20–22^. Through the coordinated regulation of their expression, PXR can directly influence the level of active metabolites of these drugs and their delivery into tumor cells.

PXR expression has been described in multiple cancer types^23–28^, where it has been implicated in the modulation of tumor cell responses to various antineoplastic drugs. For instance, in colorectal cancer, PXR promotes chemoresistance to 5-FU/irinotecan-based chemotherapy by regulating a network of genes involved in drug transport and metabolism, including ABCG2, CYP3A4 and UGT1A1^28,29^. PXR silencing, as well as its downregulation through niclosamide-induced expression of miR-148a, sensitized cells to chemotherapy *in vitro* and reduced tumor relapse *in vivo* in xenograft models^26^. Another study showed that the PXR antagonist SPA70 sensitized paclitaxel-resistant non-small cell lung cancer cells *in vitro* and reduced tumor growth *in vivo* under paclitaxel treatment by disrupting PXR binding to the ABCB1 promoter thereby reducing P-glycoprotein expression^24^. Conversely, PXR can also enhance sensitivity to other classes of drugs, such as kinase inhibitors. For instance, PXR expression increases sensitivity to afatinib in 22Rv1 prostate cancer cells *in vitro* by promoting the expression of the monocarboxylate transporter SLC16A1, resulting in increased intracellular accumulation of this inhibitor of the ErbB family of receptor tyrosine kinases^23^.

However, the role of PXR may extend beyond its metabolic functions, as it was also associated with cell proliferation^30^ and apoptosis^31,32^, and may also contribute to cancer cell plasticity as it has been identified as a specific marker of cancer stem cells in colorectal cancer^28^. These observations raise the possibility that PXR could influence therapeutic responses through other mechanisms than metabolism per se.

The potential involvement of PXR in melanoma is of particular interest in the context of targeted therapies. First, PXR expression has never been assessed in melanoma. Furthermore, the BRAF inhibitor dabrafenib that is clinically approved in the treatment of melanoma, was shown to display a strong PXR agonistic activity that was greater than that of SR12813^33^, a reference PXR agonist commonly used to study PXR at the functional level^34^. This prompted us to investigate the expression of PXR in melanoma and study its role in the response to BRAFi and MEKi. Our results show for the first time that PXR is expressed at a significant level in a large proportion of melanoma samples. They also demonstrate that PXR sensitizes BRAF-V600E mutated melanoma cell lines to BRAFi, MEKi and their combinations and that this effect is associated with a shift toward a more differentiated phenotypic state of melanoma cells, highlighting a new role for PXR that is independent of its canonical role in drug metabolism or transport.

## MATERIALS AND METHODS

### Drugs and reagents

Dabrafenib (S2807), trametinib (S2673), encorafenib (S7108), binimetinib (S7007), vemurafenib (S1267) and cobimetinib (S8041) were purchased from Selleck Chemicals. Dimethyl sulfoxide (DMSO), SPA70 (SML2662), dextran from Leuconostoc mesenteroides (D8821), activated charcoal (C9157), and Propidium Iodide (P4170) were purchased from Sigma-Aldrich. Hoechst 33342 was purchased from Fisher Scientific. RNAse A was purchased from QIAGEN. SR12813 (126411-39-0) was purchased from MedChemExpress.

### Cell lines

The BRAF-V600E-mutated melanoma cell lines A375 and SK-MEL-28 were obtained from the American Type Culture Collection (ATCC) and cultured in DMEM+GlutaMAX™ medium (Thermo Fisher Scientific) supplemented with 10% FBS, without penicillin/streptomycin at 37°C in a 5% CO2 humidified atmosphere. Cells expressing PXR and control cells were generated by electroporation of either the pcDNA3-hPXR plasmid, encoding human PXR under the control of the CMV promoter (a gift from Dr. JM Pascussi, IGF, Montpellier, France), or the corresponding empty vector, as described previously^23^. For A375 cells, co-transfection with the XREM-CYP3A4-Luc+ luciferase reporter plasmid was performed. For each electroporation, 5 x 10^6^ cells were resuspended in 500 µL Opti-MEM™ medium (Thermo Fisher Scientific) and electroporated with 10 µg of each plasmid at 250 V and 500 µF. Stable clones were isolated by limiting dilution in the presence of 700 µg/mL geneticin (G418; InvivoGen) and 175 µg/mL Hygromycin B Gold (InvivoGen) for A375 clones, and 700 µg/mL geneticin (G418; InvivoGen) for SK-MEL-28 clones. Cell lines were routinely tested for mycoplasma contamination using the MycoAlertTM detection kit (Lonza).

### Tissue Micro Arrays (TMA) and immunohistochemistry

A malignant melanoma test TMA, containing 10 cases of malignant melanoma and 2 cases of skin tissue (T382c), and a malignant melanoma TMA, containing 45 cases of malignant melanoma, 1 case of adjacent normal skin tissue, and 4 cases of skin tissue (ME1002b), were purchased from TissueArray.com LLC. In addition, formalin-fixed, paraffin-embedded (FFPE) melanoma samples from 52 patients collected before treatment with dabrafenib/trametinib were obtained from the Biological Resource Center (CRB) of Nîmes University Hospital, adhering to ethical requirements regarding non-objection to biomedical research. Immunohistochemical staining was performed on a Leica Bond Rx automated staining platform using the Bond Polymer Refine Detection kit adapted for alkaline phosphatase detection (DS9390, Leica). Antigen retrieval was performed using ER2 solution for 20 min at 100°C. PXR was detected using a mouse monoclonal antibody (clone G-11, sc-48403; Santa Cruz Biotechnology) diluted 1:100 and incubated for 15 min at room temperature, followed by incubation with the post-primary reagent and AP-conjugated polymer according to the manufacturer’s protocol. Validation of the PXR immunostaining has been reported in a previous study^23^. Signal was developed using Fast Red chromogen to allow discrimination of the immunohistochemical signal from endogenous melanin pigment, which can be difficult to distinguish from the brown precipitate generated by DAB in pigmented lesions. Sections were counterstained with hematoxylin, dehydrated, and mounted. Slides were digitized at 20× magnification using a 3D-Histech scanner (MRI Imaging Platform). Immunostaining was performed by the Translational Research Unit at the Montpellier Cancer Institute (ICM). Quantification of cytoplasmic PXR expression was performed by a pathologist using a semi-quantitative H-score, calculated as follows: H-score = Σ(i × %), where i represents staining intensity (graded from 0 to 3) and % represents the percentage of tumor cells showing each staining intensity. H-scores therefore ranged from 0 (no positive cells) to 300 (100% of cells with maximal staining intensity).

### Transactivation Assay for PXR Agonist Activity

The HG5LN and HG5LN GAL4-hPXR cell lines were used to assess the ability of xenobiotics to activate human PXR. HG5LN cells were derived from the HeLa cell line and stably transfected with the GAL4RE5-βGlob-Luc-SVNeo plasmid, resulting in expression of a luciferase reporter gene under the control of five GAL4 response elements. HG5LN-GAL4-hPXR cells were generated by co-transfection with the GAL4RE5-βGlob-Luc-SVNeo and pSG5-GAL4(DBD)-hPXR(LBD)-puro plasmids, resulting in the expression of a chimeric protein comprising the yeast GAL4 DNA-binding domain (DBD) fused to the human PXR ligand-binding domain (LBD)^35^. HG5LN and HG5LN GAL4-hPXR cells were seeded in 96-well, white, opaque, flat bottom plates at 40,000 cells per well in 150 µL of DMEM/F12 medium without phenol red supplemented with 2.5% dextran-coated charcoal-treated fetal bovine serum prepared in-house (DCC-FBS). Compounds were added 24 h later using an automated workstation (Biomek 3000, Beckman Coulter, Villepinte, Paris), and cells were incubated at 37°C for 16 h. Then, medium was removed, and 50 µL of test medium containing D-Luciferin (#12507, AAT Bioquest) at 0.3 mM was added per well. Luciferase activity was measured for 2 s in intact living cells after 10 min stabilization using a MicroBetaWallac luminometer (PerkinElmer). Each compound was tested at various concentrations in three independent experiments. Individual agonist dose-response curves were fitted using the sigmoid dose-response function of GraphPad Prism and allowed the estimation of EC_50_ (effective concentration for half-maximal luciferase activity) values.

### Western blotting

Protein lysates were performed using Laemmli buffer supplemented with dithiothreitol (DTT; 5% v/v) and 500 U/mL Benzonase^®^ (Sigma-Aldrich). Samples were heated at 95°C for 5 min. Lysates (50 µg) were electrophoresed in 10% SDS-PAGE (120 V for 60 min) and transferred onto nitrocellulose membranes using the Trans-Blot^®^ Turbo Transfer System (Bio-Rad Laboratories). Total protein staining was performed using the Revert™ 700 Total Protein Stain Kit (LICORbio) according to the manufacturer’s instructions, and the signal was acquired immediately after staining. Then, membranes were blocked with 5% milk in PBS containing 0.1% Tween 20 (PBS-T) and incubated overnight at 4°C with primary antibodies diluted in 5% BSA in PBS-T. Membranes were subsequently incubated for 1 h with IRDye^®^ secondary fluorescent antibodies (LICORbio) diluted 1/10,000 in 5% BSA in PBS-T. Protein signals were detected using an Odyssey^®^ XF imaging system (LICORbio) and quantification was performed using the Image Studio™ Software (LICORbio). The list of the antibodies used in this study is provided in the Supplementary Table 1.

### RT-qPCR

For RNA extraction, cells were seeded in 100mm Petri dishes (10^6^ cells per well) in DMEM/F12 medium without phenol red and complemented with 2.5% DCC-FBS. The day after, cells were treated with the indicated treatments. At the end of the treatments, total RNA was extracted using the Quick RNA Miniprep (Zymo Research) kit, according to the manufacturer’s instructions. RNA concentrations were measured using a Nanodrop 2000 (Thermo Scientific). Then, 500 ng of total RNA was mixed with 0.5 µL of random primers, 1 µL of dNTP mix, and nuclease-free water to a final volume of 14 µL and incubated at 65°C for 5 min and then placed on ice for at least 1 min. Subsequently, 4 µL of 5X First-Strand Buffer, 1 µL of 0.1 M dithiothreitol (DTT), and 1 µL of SuperScript™ III reverse transcriptase were added. The reaction mixture was incubated at 25°C for 5 min, 50°C for 60 min, and 70°C for 15 min. The resulting cDNA was stored at −20°C until use. Expression level of the different genes was evaluated by real-time quantitative PCR using a LightCycler 480 system in the presence of ONEGreen FAST qPCR Premix (Ozyme, Saint-Cyr-l’École, France) and DNA primers used at 10 µM (final concentration). Primers were purchased from Eurofins Genomics (Ebersberg, Germany). Sequences of the primers that were used are shown in Supplementary Table 2. Expression levels were calculated using the ΔΔCt method and were normalized to the expression of RPL0 used as a housekeeping gene.

### Cytotoxicity assays

The effect of the different drugs on cell growth was assessed using the sulforhodamine B (SRB) assay. Briefly, cells were seeded in 96-well plates in phenol red-free DMEM/F12 medium supplemented with 2.5% dextran-coated charcoal-treated fetal bovine serum (DCC-FBS). For A375 cells, E17, E30, and 1D11 clones were seeded at 2500 cells/well, 4B6 and 5E6 clones at 3000 cells/well, and pcDNA, pcDNA2, and WT clones at 2000 cells/well. All the SK-MEL-28 clones were seeded at 2500 cells/well. The following day, cells were treated with the indicated drug concentrations for 72 h. For BRAFi/MEKi combination treatments, concentration matrices were used, in which increasing concentrations of BRAFi were tested in combination with increasing concentrations of MEKi^36^. A total of 48 combinations were tested in duplicate (Fig. S2). After 72 h, the medium was removed and cells were fixed overnight with 10% trichloroacetic acid. Cells were then stained with 0.4% sulforhodamine B in 1% acetic acid, washed with 1% acetic acid, and incubated with 10 mM Tris-HCl for 10 min with gentle shaking. Absorbance was measured at 560 nm using a PHERAstar FS plate reader. Percent growth was calculated relative to untreated cells (set at 100%) and plotted as a function of drug concentration. At least three independent experiments were performed for each clone and treatment condition.

Alternatively, drug cytotoxicity was evaluated by fluorescence imaging using the Celigo^®^ imaging cytometer (MRI Imaging Platform), following Hoechst 33342 and propidium iodide (PI) staining to quantify live and dead cells, respectively. Briefly, cells were stained with 5 µg/mL Hoechst 33342 and 1 µg/mL propidium iodide for 15 min without removing the culture supernatant prior to image acquisition. Staining was then quantified using a dedicated Celigo^®^ software as described previously^37^.

For BRAFi/MEKi combination treatments, “drug interaction” analyses were performed using the BRAID (Bivariate Response to Additive Interacting Doses)^38^ model implemented in the *braidrm* package (version 0.71) in R, which uses the Hill parameters from the dose-response curves of each drug to model their combined effects. Plots were generated in R (Fig. S2).

### Cell cycle analysis

Cell cycle distribution following treatment was assessed by measuring cellular DNA content using propidium iodide (PI) staining. Adherent cells were detached with trypsin and collected together with cells present in the culture supernatant to account for apoptotic cells. Cells were washed once with PBS, and 0.5-1 × 10^6^ cells were resuspended in 500 µL PBS. Cells were fixed by adding 1.5 mL ice-cold 100% ethanol while vortexing and incubated on ice for at least 10 min. Fixed cells were then centrifuged, washed with PBS, resuspended in 500-700 µL of staining solution containing 40 µg/mL propidium iodide and 100 µg/mL RNase A in PBS and incubated for 20 min at room temperature, protected from light. Samples were then directly analyzed using the Beckman Coulter Gallios flow cytometer (MRI Imaging Platform).

### Quantification of Drug Concentrations

Intracellular and extracellular concentrations of dabrafenib and trametinib were measured using HPLC-MS/MS analysis. Cells were seeded in 100-mm Petri dishes at 5 × 10^6^ cells per dish in phenol red-free DMEM/F12 medium supplemented with 2.5% DCC-FBS. The following day, the medium was replaced with fresh medium containing fixed concentrations of dabrafenib (40 or 100 nM) or trametinib (20 or 50 nM) for 2, 4, 6, 24, or 48 h. At the end of each treatment, cells and supernatants were collected separately and processed in parallel. Samples were stored at −20°C until analysis. For the measurement of intracellular drug concentrations, cell pellets were thawed, resuspended in 500 µL of water and vortexed for 1 min. Samples were centrifuged at 15,000 × g for 15 min, and 100 µL of the supernatant was mixed to 200 µL of acetonitrile containing the internal standards (^2^H_9_-dabrafenib, ^13^C_2_,^2^H_6_-carboxy-dabrafenib, and ^2^H_4_-trametinib) and vortexed for 5 s. Following centrifugation at 15,000 × g for 15 min, 100 µL of the supernatant was transferred to an analytical vial and mixed with 100 µL of a solution containing 50% water, 50% methanol, and 0.1% formic acid. Samples were vortexed for 5 s, and 10 µL was injected in an XBridge C18 column (100 × 2.1 mm, 3.5 µm) under isocratic conditions with a mobile phase consisting of 97% (v/v) methanol and water containing 0.1% (v/v) formic acid using a Shimadzu LC-20 Prominence HPLC system (Shimadzu, Tokyo, Japan) coupled to an AB SCIEX API 2000 triple quadrupole mass spectrometer (AB SCIEX, Foster City, CA, USA).

Determination of extracellular concentrations was performed using 100 µL of thawed culture supernatant mixed with 200 µL of acetonitrile containing the internal standards and processed as described above.

### Bulk RNA sequencing

A375 cells (wild-type cells, control clones or clones stably overexpressing PXR) were seeded in 100mm Petri dishes (10^6^ cells per well) in DMEM/F12 medium without phenol red complemented with 2.5% DCC-FBS. The following day, cells were treated with vehicle (DMSO, 0.0001% v/v) or with 10 nM dabrafenib for 6 h or 24 h and RNA extraction was performed using the Quick-RNA Miniprep Kit (Zymo Research), according to the manufacturer’s instructions. Three independent biological replicates were performed for each condition. For mRNA-seq library preparation, 500 ng of total RNA in 50 µL was used for end repair, A-tailing, and adapter ligation, followed by purification. Quality assessments of the libraries were performed using a TapeStation system and libraries quantified using the Quant-iFluor™ assay. Sequencing was performed on a NovaSeq X Plus platform using the NovaSeq X Plus 25B Reagent Kit (300 cycles), generating up to 25 billion reads and 7500 Gbases per run. Bioinformatic analysis of the RNA-seq data included quality control, sequence alignment, read counting, and transcriptomic data exploration using the online QuBy platform developed by the iGenSeq Core Facility at the Paris Brain Institute (ICM), including expression analysis, principal component analysis (PCA), differential expression analysis and enrichment analysis (GSEA). Differential gene expression analysis was performed using DESeq2 with a false discovery rate (FDR) threshold of 0.05 and an absolute log2 fold change (|log2FC|) threshold of 1. This work benefited from equipment and services provided by the iGenSeq Core Facility at the Paris Brain Institute (ICM). Raw and processed data from bulk RNA-seq of each studied condition have been deposited in the NCBI Gene Expression Omnibus (GEO) under accession number GSE352675.

### Gene regulatory network analysis

RNA-seq expression data, expressed as counts per million (CPM), were analyzed using DIANE (Dashboard for the Inference and Analysis of Networks from Expression data^39^), an interactive platform that integrates differential expression analysis and gene regulatory network inference using GENIE3-based Random Forests. A minimum gene count sum of 41 across the 41 samples was used as the filtering threshold. Differentially expressed genes were identified using edgeR within the DIANE platform, with an FDR threshold of 0.05 and an absolute log2 fold change threshold of 1. The resulting list of differentially expressed genes was used as input for GENIE3-based gene regulatory network inference. A predefined list of genes was provided to DIANE as candidate regulators for gene regulatory network inference. This list comprised NR1I2, RXRG, SOX10, MITF, ZEB1, ZEB2, SNAI1, SNAI2, TWIST1, JUN, JUNB, FOS, FOB, FOSL1, FOSL2, MXI1, FOSL1, PRRX1, PRRX2, TFAP2A, TFAP2B, MEF2C, SOX6, TCF4, NFATC2, PAX3, SOX9, EGR3 and ETV4. GENIE3 uses Random Forest regression to infer the relative importance of candidate regulators for the expression of target genes, generating weighted regulator-target links. Regulators showing a correlation greater than 90% were grouped in the network inference to reduce redundancy between highly correlated predictors, as implemented in DIANE. Random Forests were constructed using 1000 trees, and the network was filtered to a connectivity density of 0.05, retaining the strongest 5% of inferred regulator-target links. The resulting networks were visualized and processed using Cytoscape for figure generation.

### Statistical Analysis

Statistical significance was assessed using the non-parametric Wilcoxon test or Kruskal-Wallis test, as appropriate. Graphs, tables, and statistical analyses were generated using GraphPad Prism and Microsoft Excel. ^∗^p value < 0.05, ^∗∗^p value < 0.01, ^∗∗∗^p value < 0.001.

## RESULTS

### PXR is expressed in melanoma and is activated by BRAF inhibitors used in the clinic

PXR expression levels have been measured in many tumor samples from various origins but were never evaluated in melanoma. Therefore, we assessed PXR expression by immunohistochemistry in tumor samples from 100 patients spanning all stages of melanoma (Fig. 1A). The results show that PXR expression was detected in the majority of melanoma samples, with only 9% of samples with no detectable expression (Fig. 1B). Although nuclear localization was observed in some cases, PXR staining was predominantly cytoplasmic (Fig. 1A). Levels of PXR expression were calculated for each sample using a semi-quantitative H-score, incorporating both staining intensity and the percentage of tumor cells exhibiting each intensity level. Among the analyzed samples, 52% exhibited low PXR expression (H-score 0-100), 25% moderate expression (H-score >100 - <200), and 14% high expression (H-score ≥ 200) (Fig. 1B and S1A), but its expression was not significantly associated with the disease stage (Fig. S1B). Since we previously showed that dabrafenib was a potent agonist of PXR^33^, we further investigated whether other BRAFi and MEKi that are used in combination for the treatment of metastatic melanoma could also modulate PXR activity. For this purpose, we used a luciferase reporter assay developed in metabolically deficient HG5LN cells stably overexpressing the GAL4-hPXR fusion protein^35^. All three BRAFi commonly used in the clinic (dabrafenib, encorafenib, and vemurafenib) exhibited PXR agonistic activity *in vitro* following 24 h treatment (Fig. 1C). Dabrafenib activated PXR at relatively low concentrations, with an estimated EC_50_ of 73.5 nM (95% CI, 54.9-108.8 nM), compared with 228.8 nM (95% CI, 180.8-294.1 nM) for the reference PXR agonist SR12813. Encorafenib and vemurafenib also activated PXR but at concentrations above 100 nM, with an estimated EC_50_ of 7.8 µM (95% CI, 6.4-10 µM) and >10 µM for encorafenib and vemurafenib, respectively (Fig. 1C and S1C). In contrast, the three MEKi that were tested (trametinib, binimetinib and cobimetinib) did not exhibit PXR agonistic activity. Though luciferase activity was induced by binimetinib and cobimetinib for concentrations above 300 nM in HG5LN GAL4-hPXR cells, it was also induced in control HG5LN cells, indicating a PXR-independent, non-specific effect (Fig. 1C). Overall, these results demonstrate that clinically used BRAFi, but not MEKi, can activate PXR *in vitro*.

**Figure 1.**
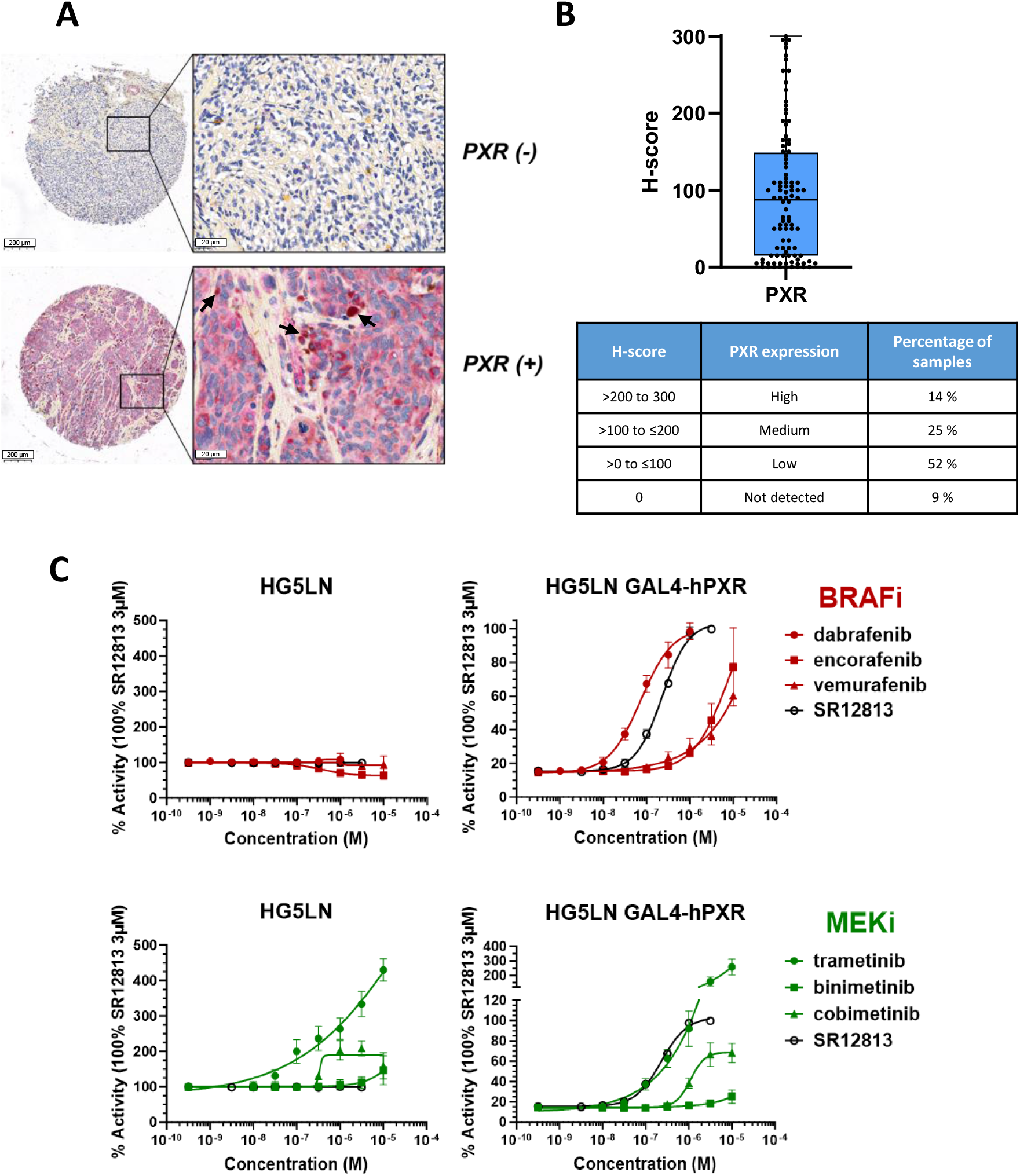
PXR expression in clinical melanoma samples and activation by BRAF inhibitors. **A)** Representative images of PXR immunohistochemical staining (red) in melanoma. Arrows show intense staining. Scale bars: 200 μm (low magnification) and 20 μm (high magnification). **B)** Distribution of PXR expression levels across 100 patient samples, quantified using the H-score method (staining intensity, ranging from 0 to 3, multiplied by the percentage of cells exhibiting the corresponding intensity). **C)** Analysis of the agonistic activity of BRAF and MEK inhibitors after 24 h of treatment using the HG5LN GAL4-hPXR luciferase reporter assay. The reference agonist SR12813 was used as a positive control.

### PXR sensitizes melanoma cells to BRAFi/MEKi combinations

Given the reported role of PXR in modulating the therapeutic response to various chemotherapies across multiple malignancies, we investigated whether PXR could influence the sensitivity of melanoma cells to BRAFi and MEKi. To this end, we generated clones stably overexpressing PXR by using the BRAF-V600E-mutant melanoma cell lines A375 and SK-MEL-28. Western blot analyses showed that PXR was not detectable in wild-type cells and in control clones transfected with the empty vector whereas clones transfected with the pcDNA3-hPXR plasmid exhibited different levels of PXR expression, recapitulating the heterogeneity of its expression observed in the clinical samples (Fig. 2A and Fig. S1A). Stable transfections did not significantly affect cell growth as doubling times of A375 clones were similar (∼30 h), and only slight differences were observed for SK-MEL-28 clones with a doubling time varying between ∼40 and 50 h, depending on the clone (Fig. 2B). The functionality of PXR in each clone was then verified by measuring the expression of a well-known PXR target gene, ABCB1, by RT-qPCR following treatment with the reference agonist SR12813 (Fig. 2C). The results showed that the expression of ABCB1 was induced by 3-to 5-fold in cells overexpressing PXR, whereas it remained unchanged in the corresponding control clones. Also, treatment with the PXR antagonist SPA70 led to reduced expression of ABCB1 or had no significant effect depending on the clone (Fig. 2C). Together, these results confirmed that PXR was functional in the different clones overexpressing this nuclear receptor.

**Figure 2.**
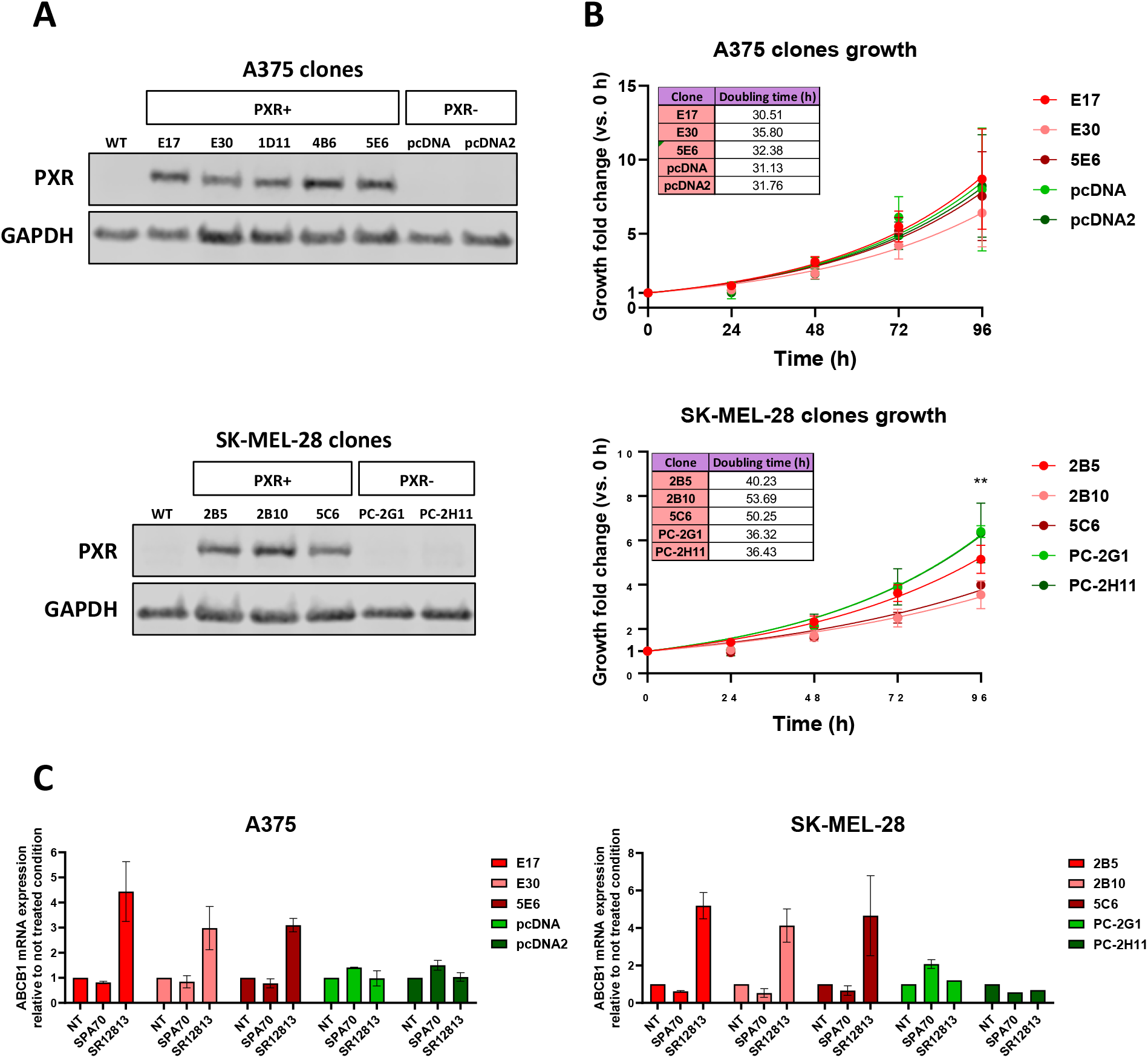
Characterization of PXR-positive (PXR+) and PXR-negative (PXR-) melanoma cell clones. **A)** Protein lysates from parental A375 and SK-MEL-28 cells (WT), as well as transfected clones from both cell lines, were prepared under basal culture conditions. PXR protein expression was assessed by Western blotting, with GAPDH used as a loading control. **B)** Growth curves of A375 and SK-MEL-28 clones under untreated conditions using the same experimental setup as the cytotoxicity assays (24 h pre-incubation followed by 72 h of treatment). Doubling times were estimated in hours. **C)** ABCB1 mRNA expression was analyzed by RT-qPCR in PXR+ and PXR-clones derived from the A375 and SK-MEL-28 cell lines. Cells were left untreated (**NT**) for 24 h or treated for 24 h with 1 μM SPA70 or 3 μM SR12813. Gene expression levels are expressed as fold changes relative to the corresponding untreated condition, which was set to 1. Statistical significance was determined using the non-parametric Kruskal-Wallis test. ^∗^p value < 0.05, ^∗∗^p value < 0.01, ^∗∗∗^p value < 0.001.

We then studied the effects of PXR expression on the sensitivity of A375 and SK-MEL-28 clones to dabrafenib or trametinib used alone (Fig. 3). In A375 cells, stable overexpression of PXR led to enhanced growth inhibition following treatment with dabrafenib, with a decrease in IC_50_ that reached a maximum of 2-fold between clone 5E6 and control clone pcDNA2 (Fig. 3A). When cells were treated concomitantly with a non-toxic concentration of the PXR antagonist SPA70 (1 µM), sensitization of A375 cells to dabrafenib was strongly reduced or abrogated as compared to control clones (Fig. 3A). Unexpectedly, we found that PXR expression also sensitized A375 cells to trametinib by ∼2-fold as compared to control clones (Fig. 3A) and showed that this effect was also abrogated by the use of SPA70 (Fig. 3A). Stabe expression of PXR also sensitized SK-MEL-28 cells to dabrafenib, as cell death evaluated by image cytometry was increased by ∼1.5-fold between clone 2B5 and control clone PC-2G1 (Fig. 3A), a difference that was also abrogated by SPA70. Conversely, no sensitization was observed in the case of trametinib (Fig. 3A). To determine whether the effect of PXR was specific to these targeted therapies, we also tested the effect of PXR expression on the sensitivity to docetaxel and cabazitaxel, two clinically used chemotherapies from the taxanes family. Conversely to BRAFi and MEKi, PXR expression induced a high level of resistance to these compounds when compared to control clones (Fig. 3B).

**Figure 3.**
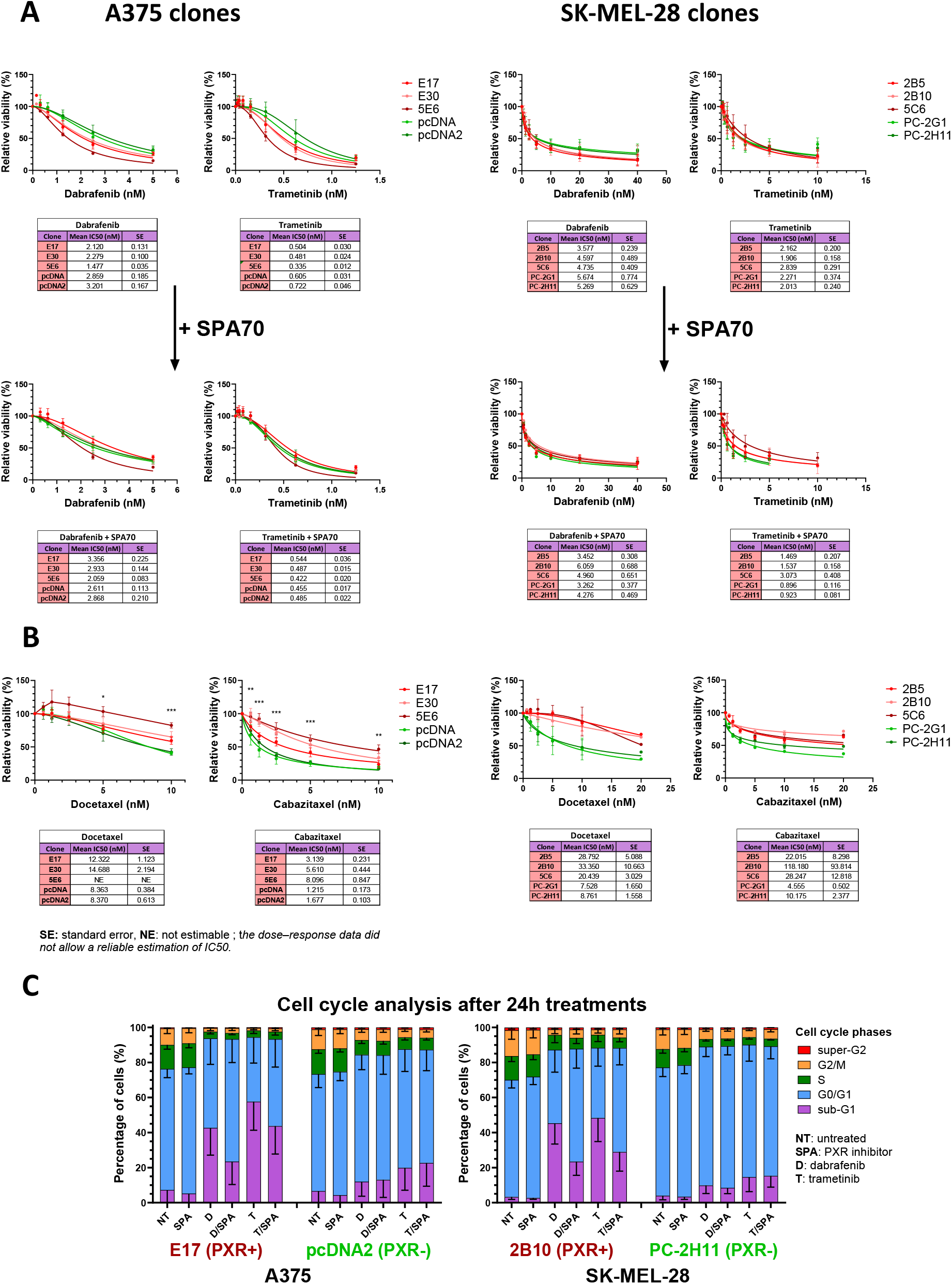
Effect of PXR expression and inhibition on the response of melanoma cells to targeted therapies and chemotherapeutic agents. **A)** Cytotoxic effects of dabrafenib and trametinib, either alone or in combination with the PXR antagonist SPA70 (1 μM), in PXR+ and PXR-clones derived from the A375 and SK-MEL-28 cell lines. Cell viability was assessed after 72 h of treatment, using the sulforhodamine B assay for the A375 clones or image analysis of propidium iodide-staining for SK-MEL-28 clones. **B)** Cytotoxic effects of docetaxel and cabazitaxel in PXR+ and PXR-clones derived from the A375 and SK-MEL-28 cell lines, assessed after 72 h of treatment, using the sulforhodamine B assay for the A375 clones or image analysis of propidium iodide-staining for SK-MEL-28 clones. **C)** DNA content measurement by propidium iodide staining after 24 h of treatment in PXR+ and PXR-clones derived from the A375 and SK-MEL-28 cell lines. Cells were treated with dabrafenib (10 nM for A375, 40 nM for SK-MEL-28) and trametinib (5 nM for A375, 10 nM for SK-MEL-28), either alone or in combination with the PXR antagonist SPA70 (1 μM). Statistical significance was determined using the non-parametric Kruskal-Wallis test. ^∗^p value < 0.05, ^∗∗^p value < 0.01, ^∗∗∗^p value < 0.001.

To further investigate the effects of BRAFi and MEKi on cell death, we performed cell cycle analysis following treatment with BRAFi or MEKi (Fig. 3C). The results showed that, in the A375 E17 clone stably expressing PXR, treatment with dabrafenib (10 nM, 24 h) and trametinib (5 nM, 24 h) induced a significant accumulation of apoptotic cells in the sub-G1 phase (∼40% and ∼55%, respectively), compared with the control pcDNA2 clone, in which sub-G1 accumulation was markedly lower (∼10% and ∼15%, respectively). When cells were treated in the presence of SPA70, accumulation in the sub-G1 phase was only partially reduced in the E17 clone (to 20% for dabrafenib and 40% for trametinib, respectively), whereas it was unchanged in the pcDNA2 control clone (Fig. 3C). Similar results were obtained for the BRAFi encorafenib and the MEKi binimetinib (Fig. S2).

Together these results indicate that the effect of PXR on BRAF-mutated melanoma cell response to treatments is drug specific and that PXR is functionally involved in their sensitization to dabrafenib or trametinib. As trametinib lacks PXR agonistic activity (Fig. 1C), they also suggest that PXR-mediated effects on cell response to BRAFi and MEKi may not require the activation of PXR by an agonist. To further confirm this hypothesis, we treated A375 clones for 72 h with GL184, a dabrafenib analog that was designed to lack PXR agonistic activity while retaining its BRAF inhibitory function (compound 4c from Schneider et al., 2022^40^) (Fig. S3A). Indeed, when compared to the activity of the reference agonist SR12813 in HG5LN GAL4-hPXR reporter cell line, GL184 could not induce PXR activation, even at micromolar concentrations (Fig. S3A). Treatment of A375 cells with GL184 showed that 1D11 and E17 clones overexpressing PXR remained hypersensitive to the BRAFi (IC_50_: 20.3 and 15.4 nM, respectively) as compared to control clones pcDNA and pcDNA2 (IC_50_: 28.2 and 31.7 nM, respectively) (Fig. S3B), confirming that activation of PXR by an agonist is not necessary for the sensitization of melanoma cells to BRAFi and MEKi.

Since BRAFi and MEKi are commonly used in combination to treat tumors with specific BRAF-V600 mutations, we studied the effects of PXR expression on A375 and SK-MEL-28 cell sensitivity to the dabrafenib/trametinib association using a dedicated matrix-based approach^36,38^ (Fig. 4A and S4). Cells were treated for 72 h with various concentrations of the combination and cell response was evaluated as described previously. In order to facilitate the visualization and the comparison of cell response to the combination between the group of control clones and the group of clones overexpressing PXR, we generated isobolograms representing the concentrations of both drugs in the combination that are required to inhibit the growth of 50% of cells (IC_50_) or to achieve 25% cell death relative to untreated clones (Fig. 4B, S5A and S5B). In A375 cells, PXR overexpression conferred a higher sensitivity to the combination of dabrafenib and trametinib as compared to control clones, confirming our previous findings with drugs used alone. Differences between clones overexpressing PXR and control clones or wild-type cells in the response to the combination were abrogated in the presence of SPA70 (Fig. 4B and S5A), further confirming that sensitization to this combination involves PXR activity. Similar results were obtained when cells were treated with the combination of encorafenib and binimetinib (Fig. 4C). In SK-MEL-28 clones, PXR overexpression also induced a sensitization to the combination of dabrafenib with trametinib as concentrations of each drug to achieve 25% cell death were significantly lower than in control clones (Fig. S5B). As in A375 cells, sensitization was attenuated when SPA70 was added to the combination and was more pronounced for dabrafenib than for trametinib (Fig. S5B). Together, these results further reinforce the functional role of PXR in the response of melanoma cells to BRAFi and MEKi combinations.

**Figure 4.**
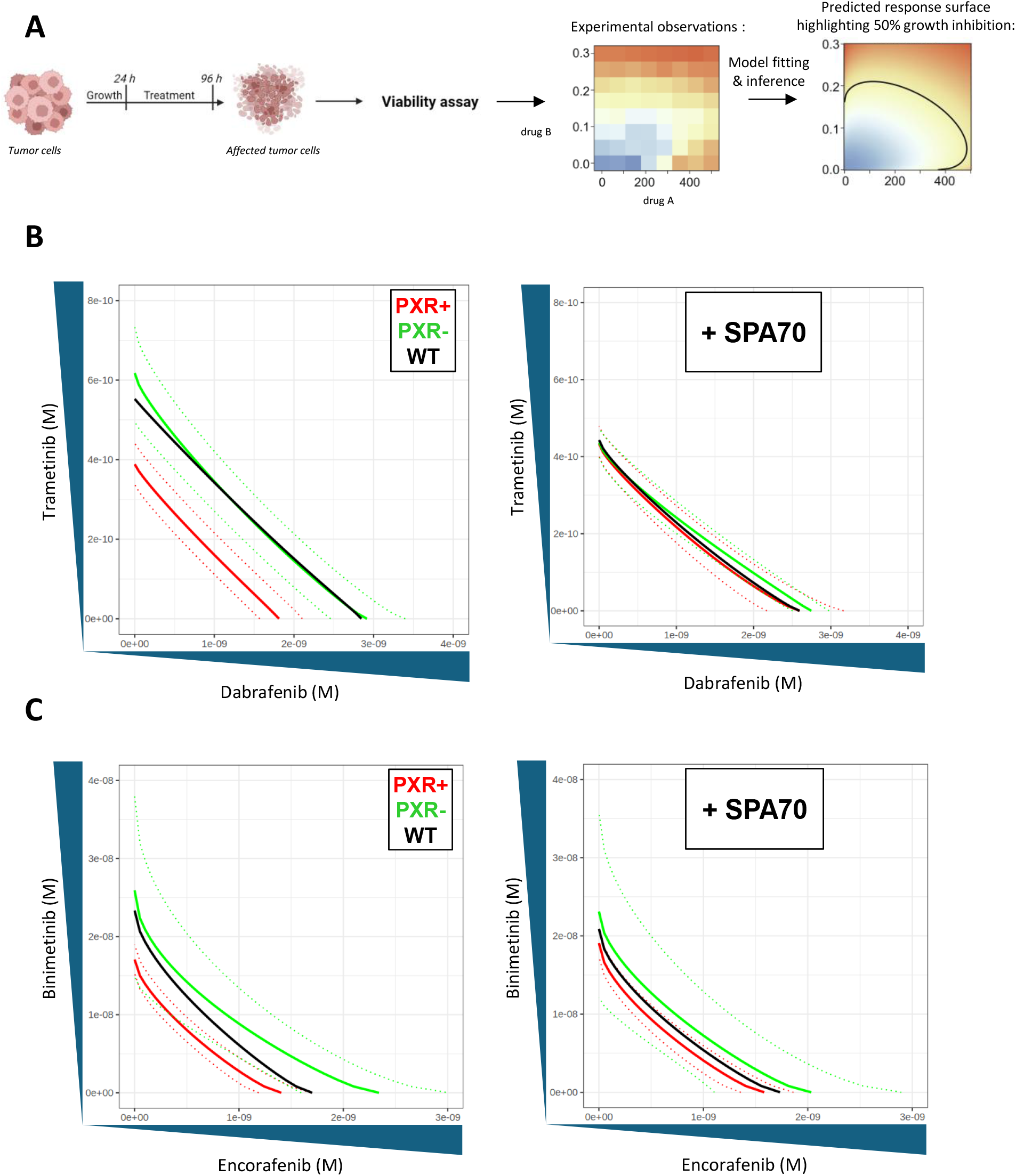
Effect of PXR expression and inhibition on the response of melanoma cells to combinations of BRAFi and MEKi. **A)** Experimental procedure used to assess clone sensitivity to drug combinations. Melanoma cells were treated in vitro with drug combinations for 72 h, after which cell viability was assessed. The isobolograms show the drug concentrations required to achieve 50% growth inhibition relative to untreated controls following combination treatment. Solid lines represent the mean responses of PXR+ and PXR-clones and parental (WT) A375 cells, while dotted lines indicate the 95% confidence intervals accounting for variability across experiments and clones. **B)** IC_50_ isobolograms showing the effect of PXR overexpression in A375 clones on relative cell viability following treatment with dabrafenib and trametinib, in the absence or presence of the PXR antagonist SPA70 (1 μM). Cell viability was assessed using the sulforhodamine B assay after 72 h of treatment. **C)** Same as in B), using encorafenib/binimetinib treatment.

### PXR does not influence the uptake of dabrafenib and trametinib during short-term treatment

To investigate the mechanism underlying PXR-mediated sensitization to BRAFi and MEKi, we first examined whether PXR could alter the intracellular accumulation or the metabolism of these targeted therapies. This possibility was supported by previous findings showing that PXR overexpression in prostate cancer cells increased the intracellular accumulation of the kinase inhibitor afatinib, thereby enhancing its efficacy *in vitro*^23^. We therefore measured the intracellular and extracellular concentrations of dabrafenib and trametinib by HPLC-MS/MS at multiple time points (Fig. 5A). The results showed that PXR overexpression was not associated with significant differences in the relative intracellular concentration of dabrafenib or trametinib between E17, 5E6 and pcDNA2 clones over time in A375 cells (Fig. 5B and S6). These results suggest that PXR-mediated sensitization to dabrafenib and trametinib is not due to altered intracellular accumulation of these kinase inhibitors.

**Figure 5.**
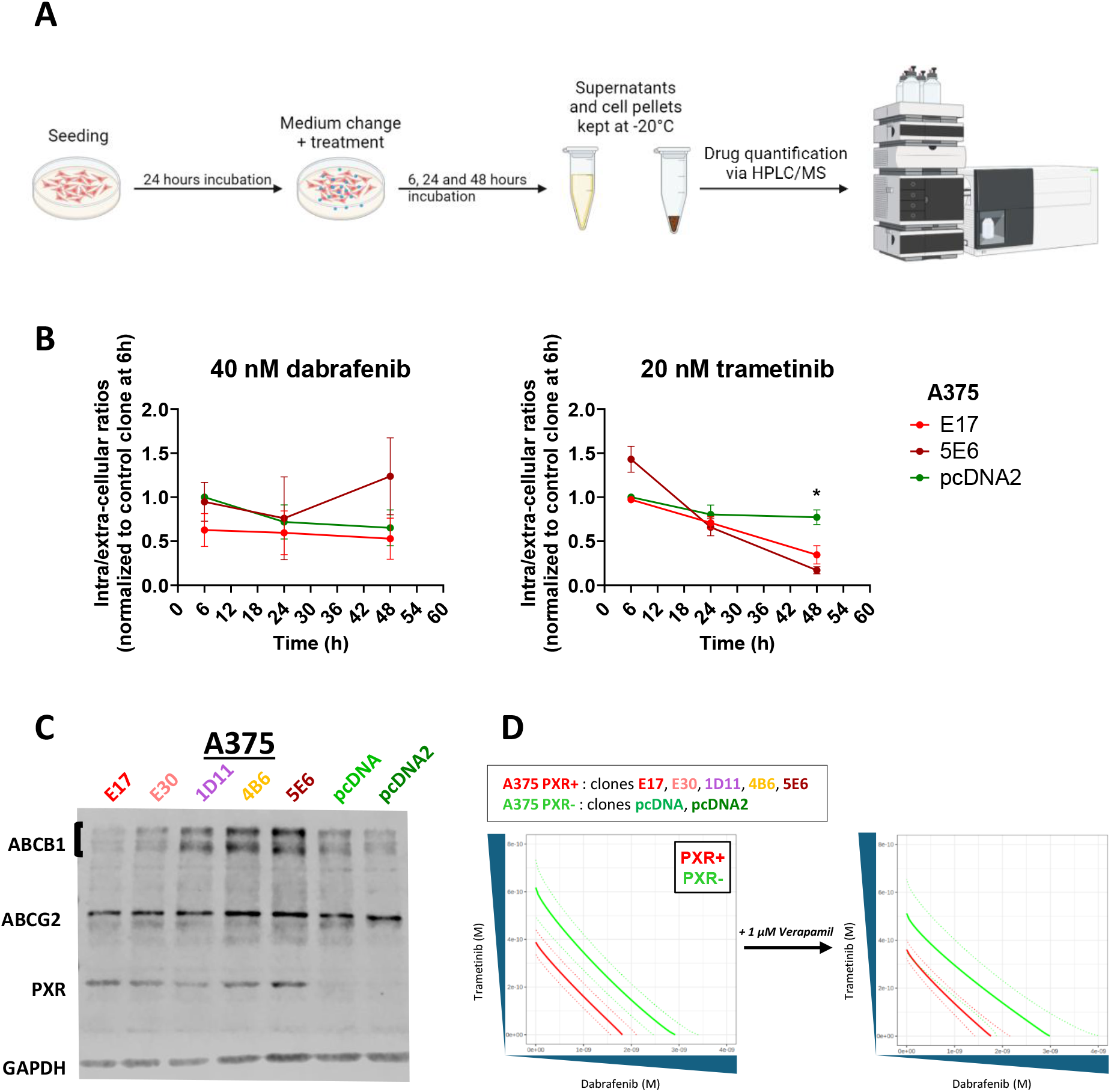
Effect of stable PXR expression on the intracellular accumulation of dabrafenib and trametinib. **A)** Experimental procedure used to assess the intracellular accumulation of dabrafenib and trametinib. **B)** A375 clones stably expressing PXR (E17 and 5E6) and the control clone (pcDNA2) were treated with 40 nM dabrafenib or 20 nM trametinib for 6, 24, and 48 h. Drug concentrations were quantified by mass spectrometry, as described in the Materials and Methods section. Results are expressed as the intracellular/extracellular ratio of drug concentrations, normalized to the earliest time point of the control clone (pcDNA2), which was set to 1. Data are presented as mean ± SD from at least two independent experiments. **C)** Protein lysates from A375 clones were prepared after 24 h of incubation without treatment. Protein expression levels of ABCB1, ABCG2 and PXR were assessed by Western blotting, with GAPDH used as loading control. **D)** IC_50_ isobolograms showing the effect of the ABCB1 inhibitor verapamil (1 μM) on the relative cell viability in A375 clones during concomitant treatment with dabrafenib and trametinib, as assessed by the sulforhodamine B assay. Isobolograms without verapamil are identical to those shown in figure 4B without SPA70. Statistical significance was determined using the non-parametric Kruskal-Wallis test. ^∗^p value < 0.05, ^∗∗^p value < 0.01, ^∗∗∗^p value < 0.001.

As PXR is known to regulate the expression of a wide range of genes involved in drug transport, we further assessed whether the well-known PXR target genes ABCB1^18^ encoding the P-glycoprotein efflux pump^41^, could contribute to the difference in the sensitivity of A375 clones to dabrafenib/trametinib treatment. Western blots performed in the different clones of A375 cells showed that ABCB1 expression levels were not correlated with the levels of PXR expression in untreated conditions (Fig. 5C). Similarly, ABCG2, another PXR target gene encoding an efflux transporter, was expressed at a similar level in both PXR-overexpressing and control clones (Fig. 5C). Also, despite similar levels of ABCB1 and ABCG2 expression, the E30 clone overexpressing PXR was much more sensitive to dabrafenib and trametinib than the pcDNA control clone (Fig. 3A). Furthermore, when cells were treated with the combination of dabrafenib with trametinib in the presence of verapamil, a clinically used calcium channel blocker and ABCB1 inhibitor, no significant change in drug sensitivity was observed between clones expressing PXR and control clones (Fig. 5D). Together, these results suggest that ABCB1 or ABCG2-mediated drug efflux does not account for the increased sensitivity of A375 clones stably expressing PXR to BRAFi/MEKi treatment.

### PXR expression is accompanied by a shift toward the melanocytic state

Given that PXR-mediated sensitization did not appear to involve changes in the intracellular accumulation or efflux of targeted therapies, we investigated whether PXR overexpression could involve other mechanisms through transcriptional rewiring. To this end, bulk RNA sequencing was performed on wild-type cells, control clones (pcDNA, pcDNA2) and PXR-expressing A375 clones (E17, E30, 5E6) under basal conditions as well as following 6 or 24 h of treatment with 10 nM dabrafenib (in clones pcDNA, pcDNA2, E17, E30). Principal component analysis revealed distinct transcriptomic profiles among these clones under basal conditions that followed a gradient consistent with their levels of PXR expression (Fig. 6A and 6B). The E30 clone, which expresses the lower level of PXR exhibited an intermediate transcriptomic profile when compared to the control clones and the E17 and 5E6 clones expressing the highest level of PXR (Fig. 6A and 6B). This transcriptional gradient was maintained following dabrafenib treatment (Fig. 6A), suggesting that the transcriptional state of the cells is closely associated with PXR expression and may influence their adaptation to dabrafenib treatment. To investigate the biological processes associated with these transcriptional changes, we performed Gene Set Enrichment Analysis (GSEA) comparing the transcriptomes of untreated PXR-overexpressing clones with those of control clones (Fig. 6C and S7A). The results show that the hallmark epithelial-mesenchymal transition (EMT) gene set was significantly enriched in clones stably expressing PXR (Fig. 6C), and this enrichment was maintained following dabrafenib treatment (Fig. S7A). By contrast, transcriptional signatures associated with metastatic formation and poor patient prognosis, reported by Winnepenninckx et al., 2006^42^ and Kauffmann et al., 2008^43^, were negatively enriched in these clones (Fig. S7B).

**Figure 6.**
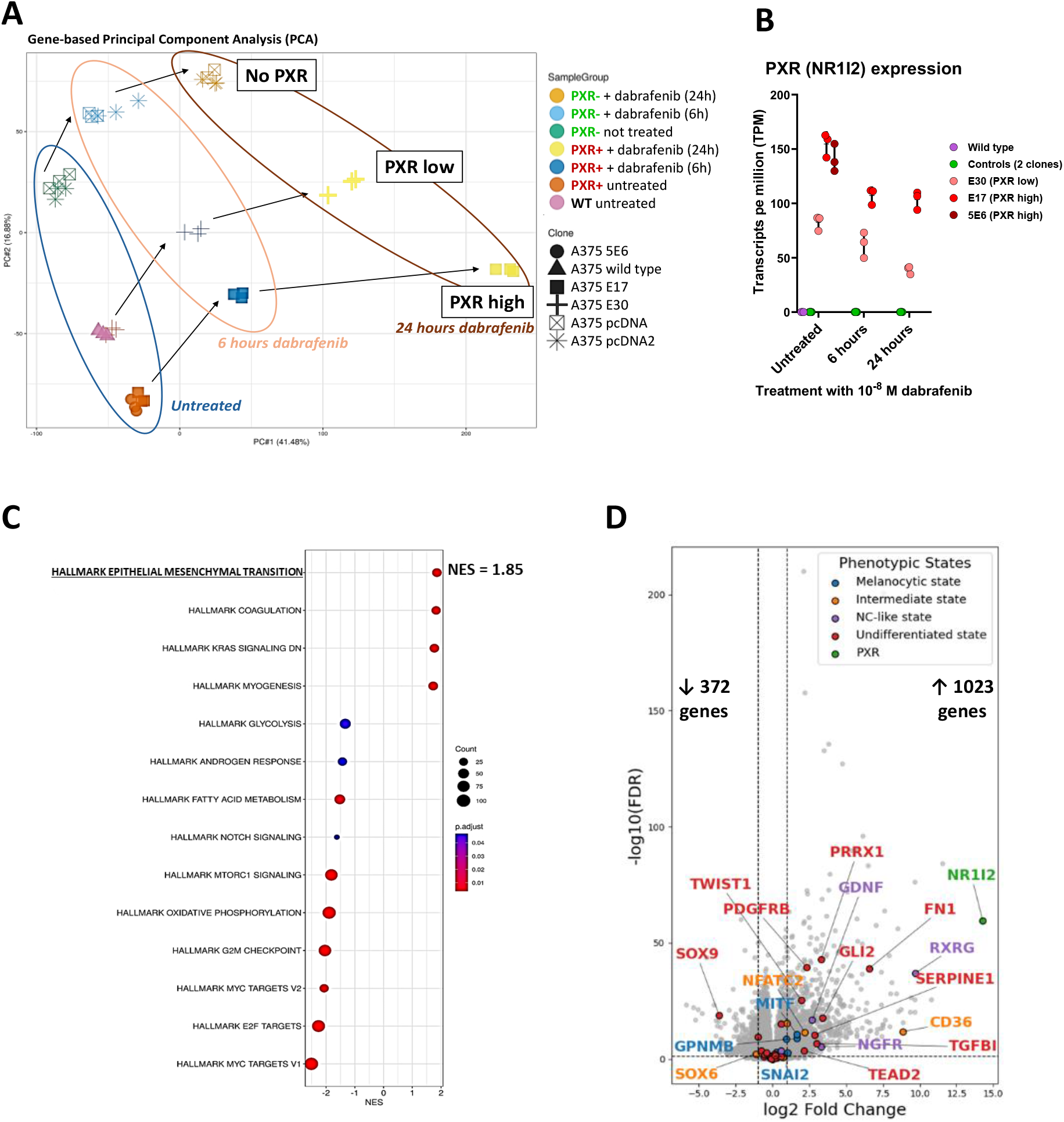
Transcriptomic evidence of a PXR-associated phenotypic shift in melanoma cells. **A)** Principal component analysis (PCA) of the transcriptomes of A375 clones under basal conditions or following 10 nM dabrafenib treatment for 6 and 24 h. Annotations were made according to treatment condition and PXR expression level. **B)** Normalized mRNA expression of PXR (NR1I2) from the RNA-seq dataset, expressed as transcripts per million (TPM). **C)** Gene Set Enrichment Analysis (GSEA) of Hallmark gene sets comparing PXR+ clones with control clones under basal conditions. Results are ranked according to the normalized enrichment score (NES), and all are statistically significant, with an adjusted p-value < 0.05. **D)** Volcano plot of differentially expressed melanoma phenotypic markers comparing PXR+ clones with control clones under basal conditions. Phenotypic markers, as defined by Pedri et al., 2022^13^, are color-coded according to their respective phenotypic state: melanocytic (blue), intermediate (orange), neural crest-like (NC-like, purple) and undifferentiated (red). NR1I2, the gene encoding PXR, is shown in green. Other genes are shown in gray. Numbers indicate the genes significantly upregulated (↑) or downregulated (↓) in PXR+ clones (|log2FC| ≥ 1 and FDR ≤ 0.05).

We next focused our study on key genes associated with melanoma phenotypic states and EMT-like processes previously reported by Pedri et al., 2022^13^ that were present among the 1395 genes differentially expressed following PXR expression in untreated conditions (Fig. 6D). These genes encode transcriptional regulators, cell-surface receptors, extracellular matrix components, and factors involved in signaling and metabolic adaptation, collectively contributing to the maintenance and plasticity of different phenotypic states. Several of these genes were significantly upregulated, including markers of the melanocytic state (MITF, GPNMB, SNAI2), of the intermediate state (NFATC2, CD36), of the neural crest-like state (RXRG, GDNF, NGFR), and of the undifferentiated state (PRRX1, FN1, SERPINE1, TGFBI, GLI2, PDGFRB, TWIST1, TEAD2) (Fig. 6D). In contrast, the transcription factors SOX9, a driver of the undifferentiated state^13^, and SOX6, a marker of the intermediate state^44^, were significantly downregulated (Fig. 6D). Notably, the fatty acid translocase CD36, which has been reported to contribute to metabolic adaptation^45^, has previously been identified as a direct transcriptional target of PXR^46^. Following dabrafenib treatment, additional melanocytic markers emerged, including ZEB2, DCT, CAPN3, TFAP2A, and IRF4, whereas fewer or no additional markers associated with the other phenotypic states were observed (Fig. S8A). Overall, an increase in melanocytic markers, including DCT, ZEB2, and GPNMB, and a decrease in invasive markers, including FN1, PRRX1, and TWIST1, were observed following dabrafenib treatment. In contrast, an increase in TGFBI, a component of the extracellular matrix that contributes to metastatic outgrowth^47^ and is associated with an undifferentiated phenotype^44^, was also observed following dabrafenib treatment (Fig. S8A, S9A and S9B).

To identify the main regulators underlying these transcriptional differences, we performed gene regulatory network analysis using GENIE3, a machine learning method that infers putative regulatory links between input regulators and target genes^39,48^. The analysis was performed on differentially expressed genes in clones overexpressing PXR. MITF emerged as a prominent regulator in the network under basal conditions, with strong regulatory links compared with the other analyzed regulators (Fig. 7A). Given its established role as a master regulator of the melanocytic state^12,49,50^, the prominent position of MITF suggests a transcriptional program associated with melanocytic differentiation. Following dabrafenib treatment, ZEB2 emerged as a prominent regulator after 6 h, followed by JUN after 24 h (Fig. S8B).

**Figure 7.**
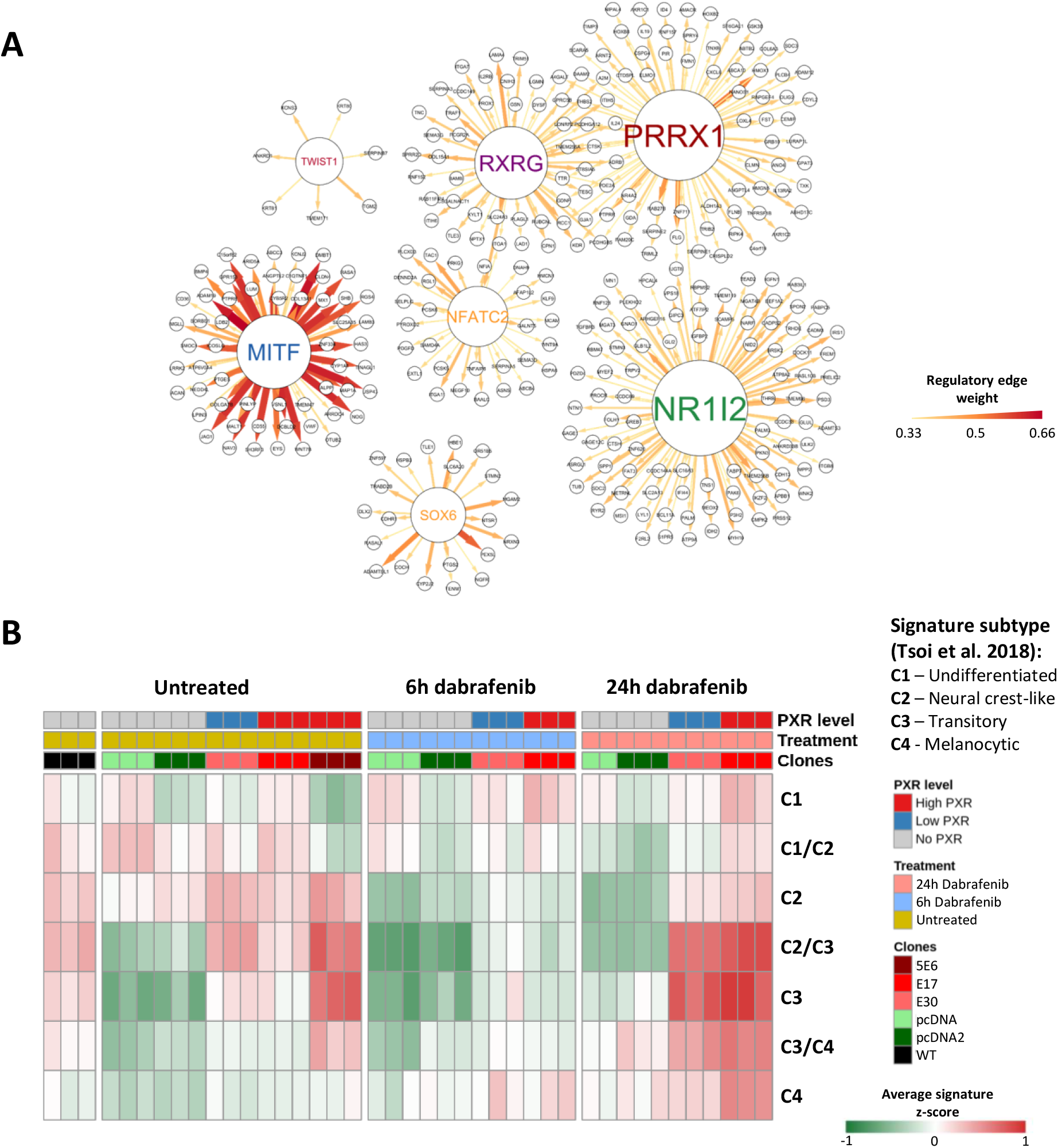
Gene regulatory network and phenotypic characterization of the PXR-associated melanoma cell state. **A)** Gene regulatory network inference analysis using GENIE3 to predict interactions among the major transcription factors involved in the melanoma phenotypic switch that were significantly dysregulated by PXR overexpression under basal conditions. Node size is proportional to the number of edges. The color intensity and thickness of the edges represent the predictive strength of each interaction and indicate the extent to which variation in the regulator predicts variation in the target gene. **B)** Heatmap of signature genes associated with the phenotypic states defined by Tsoi et al., 2018^11^, based on the transcriptomic profiles of A375 clones. The heatmap displays signature z-scores for each sample, with scaling performed independently for each signature.

Since melanoma cells are highly plastic and because their phenotypes are associated with different sensitivities to targeted therapies, we applied the phenotypic signatures defined by Tsoi et al., 2018^11^ to our bulk RNA-seq data to characterize the phenotypic changes associated with PXR overexpression under basal conditions and following dabrafenib treatment. In untreated conditions, PXR expression was associated with a marked increase in the neural crest-like/transitory hybrid state (C2/C3) and transitory state (C3) signatures as compared to control clones, particularly in the 5E6 clone (Fig. 7B and S10), indicating a PXR-mediated shift toward the transitory phenotype. This difference was less pronounced after 6 h of dabrafenib treatment, with the E17 and E30 clones showing moderately higher scores than control clones for these two signatures, with only small differences for the other signatures. After 24 h, however, both E17 and E30 clones showed increased C2/C3 and C3 signatures, while the E17 clone also showed an increase in the melanocytic state (C4) signature (Fig. 7B and S10). These results suggest that PXR expression promotes a shift toward more differentiated phenotypic states, which are associated with a higher sensitivity to BRAFi and MEKi^13^.

A375 cells display a less differentiated phenotype than SK-MEL-28 cells, which exhibit higher MITF expression^51,52^. Consistent with this difference, A375 cells have been associated with a neural crest-like transcriptional signature, whereas SK-MEL-28 cells exhibit a melanocytic transcriptional program^44^. These differences prompted us to investigate whether PXR-mediated transcriptomic rewiring could be influenced by the initial phenotypic state of the melanoma cells. We therefore analyzed the clones derived from the SK-MEL-28 cell line. In this context, PXR expression was associated with higher expression of several melanocytic markers, including MITF, DCT, and ZEB2, both in untreated cells and cells treated with dabrafenib (Fig. 8A). The levels of these markers remained consistently higher when PXR was stably expressed. PXR expression was also associated with a strong induction of E-cadherin mRNA under basal conditions, reaching a 102-fold increase in the 2B10 clone, compared with 31-fold and 10-fold increases in the 5C6 and 2B5 clones, respectively (Fig. 8A). E-cadherin expression decreased following dabrafenib treatment in clones expressing PXR, whereas it remained stable in control clones (Fig. 8A). Among markers associated with less differentiated states, only TWIST1 was upregulated in SK-MEL-28 clones expressing PXR, whether cells were treated or not, consistent with what was observed in A375 cells (Fig. S11).

**Figure 8.**
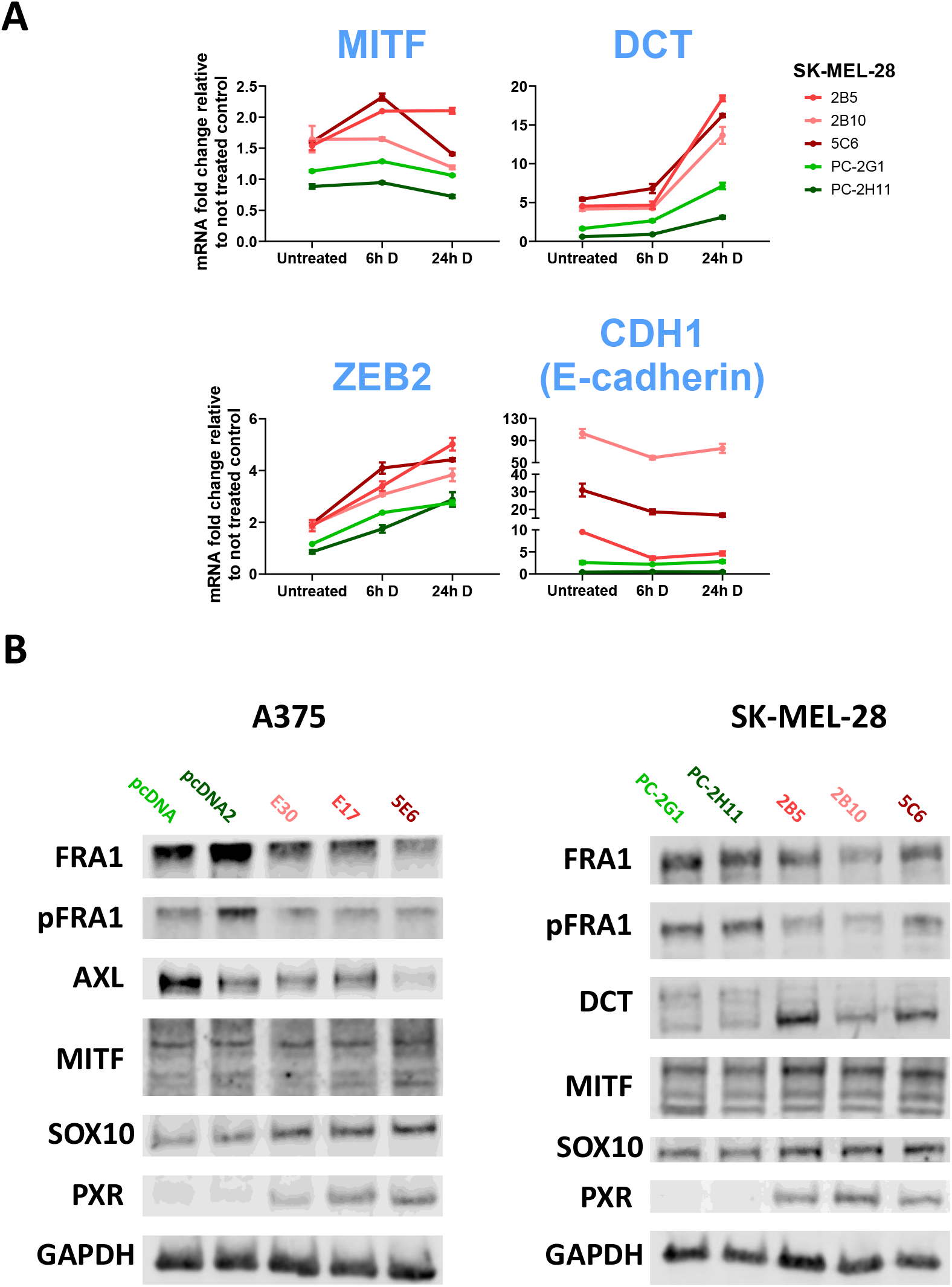
Validation of the PXR-associated phenotypic shift at the transcriptional and protein levels. **A)** RT-qPCR analysis of the expression of markers associated with the melanocytic differentiated state in untreated SK-MEL-28 clones and following treatment with 50 nM dabrafenib for 6 or 24 h. Data are presented as mean fold change relative to the average expression of the corresponding untreated controls ± SEM from one biological replicate. **B)** Protein lysates from A375 and SK-MEL-28 clones were prepared after 24 h of incubation without treatment. Protein expression levels of FRA1, phospho-FRA1 (pFRA1), AXL, DCT, MITF, SOX10, and PXR were assessed by Western blotting, with GAPDH used as control.

At the protein level, PXR expression in A375 cells was associated with a non-significant increase in SOX10 and MITF expression and a significant decrease in AXL and FRA1 expression, whereas FRA1 phosphorylation, normalized to total FRA1, was not affected (Fig. 8B and S12B). These findings were consistent with the corresponding mRNA expression data obtained in the A375 clones (Fig. S8C). Notably, the A375 5E6 clone showed the highest MITF expression, together with the lowest AXL and FRA1 levels (Fig. 8B and S12B), and exhibited the greatest sensitivity to MAPK inhibition (Fig. 3A). Similar results were obtained in SK-MEL-28 cells, in which PXR expression was associated with a significant increase in SOX10 and DCT expression and a non-significant increase in MITF expression (Fig. 8B and S13B). PXR expression also significantly reduced FRA1 phosphorylation when normalized to total FRA1, whereas total FRA1 expression was not significantly affected (Fig. 8B and S13B). Overall, these findings support the notion that PXR expression in BRAF-V600-mutant melanoma is associated with transcriptomic rewiring toward a more differentiated/melanocytic state, which is generally more sensitive to BRAF inhibition, whereas an undifferentiated state is associated with intrinsic or adaptive resistance to these targeted therapies.

## DISCUSSION

In advanced BRAF-mutant melanoma, the choice of first-line therapy is particularly important as a substantial proportion of patients may suffer disease progression despite the treatment. Immune checkpoint inhibitors are generally preferred in the first-line setting because of the potential for durable responses in a subset of patients^3,53,54^. However, BRAF/MEK inhibitor combinations can achieve faster and higher initial response rates and provide long-term responses in a subset of patients^53,54^. Beyond clinical parameters and LDH levels, which are currently used to guide treatment selection, no validated predictive biomarker is available to identify patients most likely to benefit from either strategy. In this study, we provide experimental evidences showing that PXR may be considered as a potential biomarker of sensitivity to targeted therapy in BRAF-mutant melanoma.

We first evaluated the expression of PXR in melanoma patients and show for the first time that it is detected in the majority of samples that we analyzed. Up to 39% of melanoma patients show medium to high PXR expression according to our H-score. Interestingly, although nuclear localization of PXR was observed in some cells, PXR staining was predominantly cytoplasmic. This is consistent with previous reports showing that in breast cancer positive samples, subcellular distribution of PXR was mainly cytoplasmic and only occasionally nuclear^27,55^. PXR expression located in the cytoplasm and perinuclear regions was also reported in highly differentiated tissues of non-small cell lung cancers^56^. Together, these data indicate that intracellular localization of PXR may be a dynamic process that is regulated by its co-factors and the presence of agonists.

PXR expression is known to confer a resistance to a variety of chemotherapeutic agents such as platinum derivatives, alkylating agents, DNA topoisomerase inhibitors or microtubule poisons^14,57,58^. This resistance was mainly attributable to PXR-mediated transcriptional induction of drug-metabolizing enzymes and efflux transporters, such as CYP3A4 and ABCB1 for taxanes^59,60^, the glucuronoconjugation enzyme UGT1A1 for irinotecan^29^, or the ABCC3/MRP3 transporter for oxaliplatin^61^. In this study, we confirmed that PXR expression in BRAF-V600E mutated melanoma cell lines (A375 and SK-MEL-28) also conferred a high level of resistance to taxanes while it sensitized cells to targeted therapies inhibiting BRAF and MEK.

Because we found that BRAFi, but not MEKi, could activate PXR, with dabrafenib exhibiting the strongest agonist activity^33^ (Fig. 1C and S1C), we hypothesized that this sensitization may directly result from PXR activation. However, PXR-mediated sensitization to MEK inhibitors and the fact that abrogation of the cytotoxic effects of these targeted therapies by SPA70 was only partial, demonstrate that PXR expression but not its activation by a ligand is required for the sensitization to BRAFi and MEKi. Indeed, it was previously shown that high levels of PXR expression could sustain its transcriptional activity independently of its agonists^33,62^. Similar results were reported in 22Rv1 prostate cancer cells, in which PXR expression induced a sensitization to erlotinib, dabrafenib and afatinib, whereas no effect could be observed for trametinib and vemurafenib^23^. In the case of afatinib, PXR-mediated sensitization to afatinib was due to an increase in the expression of the influx transporter SLC16A1, resulting in increased intracellular accumulation of the inhibitor^23^. In our study PXR expression had no effect on the intracellular accumulation of dabrafenib or trametinib. These data suggest that PXR-mediated effects are not only dependent on the cellular context, but also on the chemical nature of the drug. They also suggest that PXR-mediated sensitization may involve target genes other than drug transporters or metabolic enzymes, or may involve mechanisms independent of its transcriptional activity.

Besides drug pharmacokinetics, intrinsic sensitivity to MAPK pathway inhibition is influenced by both genetic alterations and phenotypic plasticity^6^. Genetic alterations such as NRAS mutations, BRAF amplification or splice variants, and MEK1/2 mutations can either reactivate the MAPK pathway or activate alternative pathways such as the PI3K-AKT pathway in order to contribute to acquired resistance^63^. In parallel, melanoma cells can switch between phenotypic states that are largely regulated by MITF^64–66^. A common distinction is based on MITF and SOX10 expression and the MITF/AXL ratio, with MITF/SOX10-low, AXL-high cells exhibiting intrinsic resistance to MAPK inhibition^12,64,65^. Other transcription factors, including FRA1, also contribute to this plasticity by promoting AXL expression and repressing MITF, thereby favoring a less differentiated and more invasive state^52^. Interestingly, in PXR-overexpressing A375 cells, we observed the upregulation of several markers associated with phenotypic plasticity, together with evidence of activation of the MITF transcriptional program. Similar findings were observed in SK-MEL-28 cells that exhibit a differentiated phenotype. Indeed, PXR expression in SK-MEL-28 cells induced the expression of several differentiation markers, including MITF. These findings are particularly relevant given the established role of MITF as a master regulator of melanocytic differentiation and melanoma cell-state plasticity^67^. Increased MITF activity can promote the expression of melanocytic differentiation markers, including TYR, TYRP1, DCT and PMEL, whereas reduced MITF activity is associated with more stem-like and tumorigenic melanoma states^50^. The initial phenotype may then directly influence the short-term response to dabrafenib in clones expressing PXR, that may undergo further differentiation during treatment, an effect that was more apparent over time based on the phenotypic signatures reported by Tsoi et al., 2018^11^. At the regulatory level, ZEB2 emerged as a prominent regulator after 6 hours following treatment and remained prominent after 24 hours, while JUN emerged at 24 hours. Interestingly, JUN upregulation is a common response to BRAFi treatment *in vitro* and in patients and has been associated with an early persistent state in melanoma cells^68^. This observation may appear at odds with the concomitant increase in the melanocytic signature that was observed following short-term dabrafenib treatment, indicative of a more differentiated phenotype. Whether these results reflect the coexistence of distinct phenotypic states within the PXR-expressing cell population or a transient adaptive state remains to be determined.

Together with our findings, these data raise the possibility that PXR expression contributes to melanoma cell-state regulation and increased sensitivity to short-term MAPK inhibition via the modulation of MITF-dependent transcriptional programs^13^, but may also trigger a long-term acquisition of resistance to MAPK pathway inhibitors in surviving cells.

PXR’s involvement in phenotypic plasticity may have broader implications for tumor progression and sensitivity to immune checkpoint inhibitors. The increased expression of melanocytic markers such as MITF and SOX10, together with decreased expression of markers of the undifferentiated state, such as AXL and FRA1, suggests that clones expressing PXR exhibit a more differentiated phenotype. This is consistent with the negative enrichment of transcriptional signatures associated with metastasis formation and poor prognosis in melanoma patients in our RNA-seq dataset, including those described by Winnepenninckx et al., 2006^42^ and by Kauffmann et al., 2008^43^.

A limitation of our study is the absence of endogenous PXR expression in the parental A375 and SK-MEL-28 cell lines, which required the generation of PXR-overexpressing clones to investigate the functional role of PXR in melanoma cells. Although our IHC analyses showed significant levels of PXR expression in a substantial proportion of melanoma patients’ samples, we were unable to detect the endogenous expression of PXR in these two BRAF V600E-mutated cell lines. This is not uncommon as the majority of cancer cell lines in culture do not express detectable levels of PXR, probably due to a transcriptional repression of this gene via the methylation of its promoter^69^. Therefore, our models may not fully recapitulate the physiological regulation of PXR observed in tumors. Additionally, the distinct phenotypic state and genetic backgrounds of SK-MEL-28 and A375 cells may have different influences on PXR-mediated effects. The sensitization to targeted therapies appeared less pronounced in SK-MEL-28 cells than in A375, which may be related to the relatively slower growth of clones expressing PXR since the cytostatic effect of MAPK inhibitors depends on cell proliferation. However, the cell death analyses by imaging of propidium iodide-stained cells and cell cycle data, which are expressed as proportions within each sample rather than relative to untreated cells, did show an increased sensitivity to BRAFi and MEKi of SK-MEL-28 clones expressing PXR, which is consistent with what was observed in A375 clones. Additional studies using models with endogenous PXR expression, ideally derived from PXR-positive patient tumors, will be useful to validate these findings.

Overall, our data provide experimental evidences that PXR can reshape the transcriptional state of melanoma cells and reinforce the notion that PXR or/and PXR-associated transcriptional signature that will need further validation could be used as a new biomarker of response to targeted therapies in BRAF-mutant melanoma.

## CONCLUSION

Our study identified PXR as a potential determinant of sensitivity to BRAFi/MEKi and their combinations in BRAF-V600E-mutant melanoma cell lines. We demonstrated that PXR expression could trigger a transcriptional program that drives melanoma cell differentiation toward a phenotype associated with increased sensitivity to MAPK pathway inhibition that did not depend on drug metabolism or transport. These findings suggest that PXR may influence not only the initial response to these targeted therapies, but may also lead to a phenotypic adaptation to the treatment and the subsequent development of resistance. Therefore, PXR expression and its associated transcriptional signature may represent promising predictive biomarkers for the identification of patients most likely to benefit from BRAFi/MEKi combinations.

## AUTHOR CONTRIBUTIONS

Conceived and designed the experiments: S.N., E.F., P.B., C.G., P.P. and A.E.

Performed the experiments: S.N., A.M., M.G., N.V., E.F., J.T. and L.T.

Analyzed the data: S.N., R.R. and M.R.

Contributed reagents/materials/analysis tools: R.R., M.R., E.F., A.B., F.L., L.M. and P.B.

Writing-original draft preparation: S.N., P.P. and A.E.

Supervised the project: P.P. and A.E.

All authors have read and agreed to the published version of the manuscript.

## SUPPLEMENTARY MATERIAL

**Figure S1.**
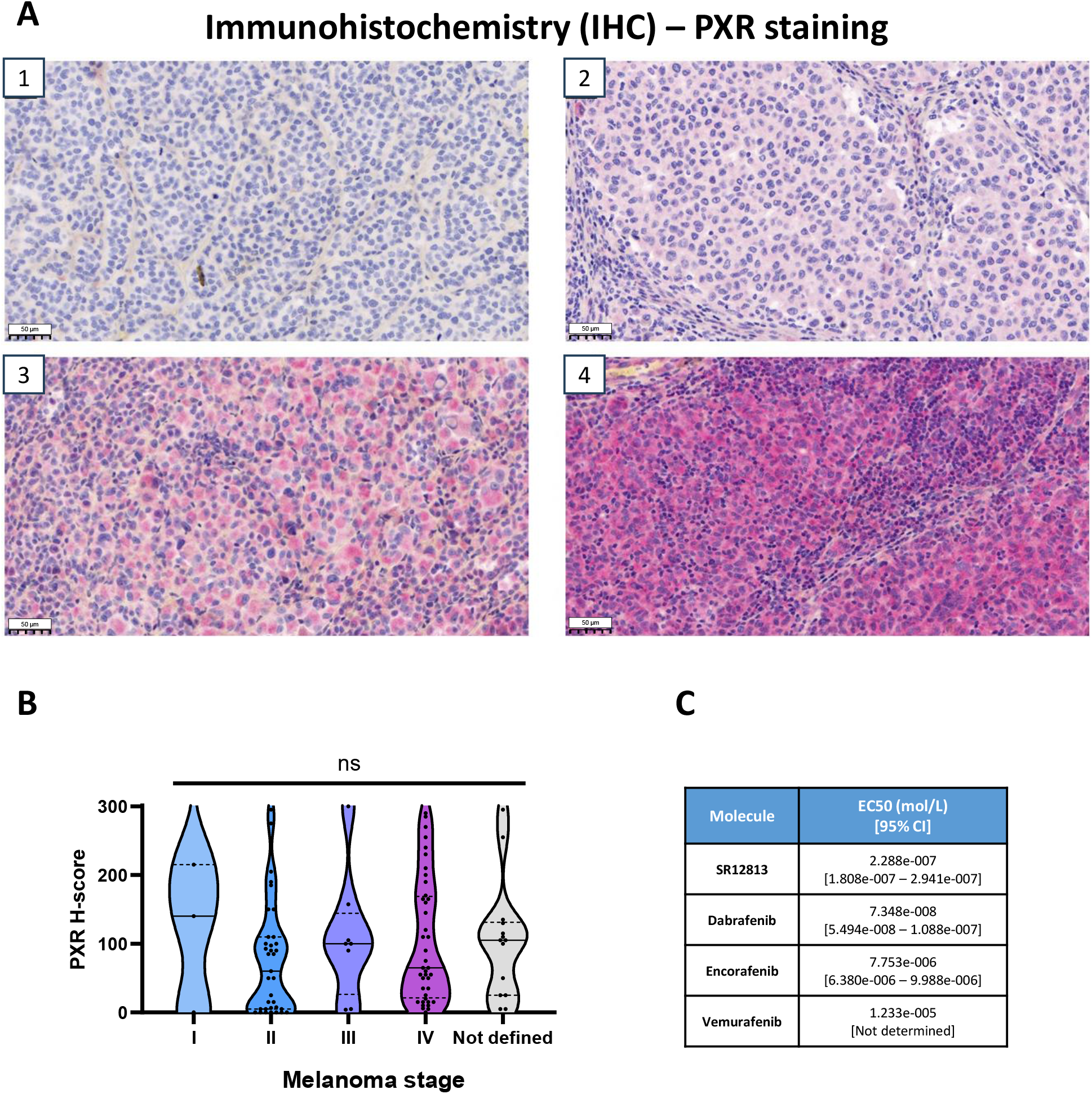
PXR expression in clinical melanoma samples and agonist activity parameters for BRAF inhibitors. **A)** PXR expression in four representative melanoma samples showing different staining intensities. **1**: No staining (score 0), **2**: Weak staining (score 1), **3**: Moderate staining (score 2), with a few tumor cells showing stronger staining (score 3). **4**: Strong staining (score 3). Cell staining scores were used to calculate the H-scores. Scale bar: 50 μm. **B)** PXR expression (H-score) across melanoma samples grouped according to AJCC stage. **C)** Estimated EC_50_ values for PXR agonism in the HG5LN GAL4-hPXR reporter model, with associated 95% confidence intervals, following nonlinear regression analysis of SR12813 and BRAF inhibitor dose-response curves. Statistical significance was determined using the non-parametric Kruskal-Wallis test. ^∗^p value < 0.05, ^∗∗^p value < 0.01, ^∗∗∗^p value < 0.001.

**Figure S2.**
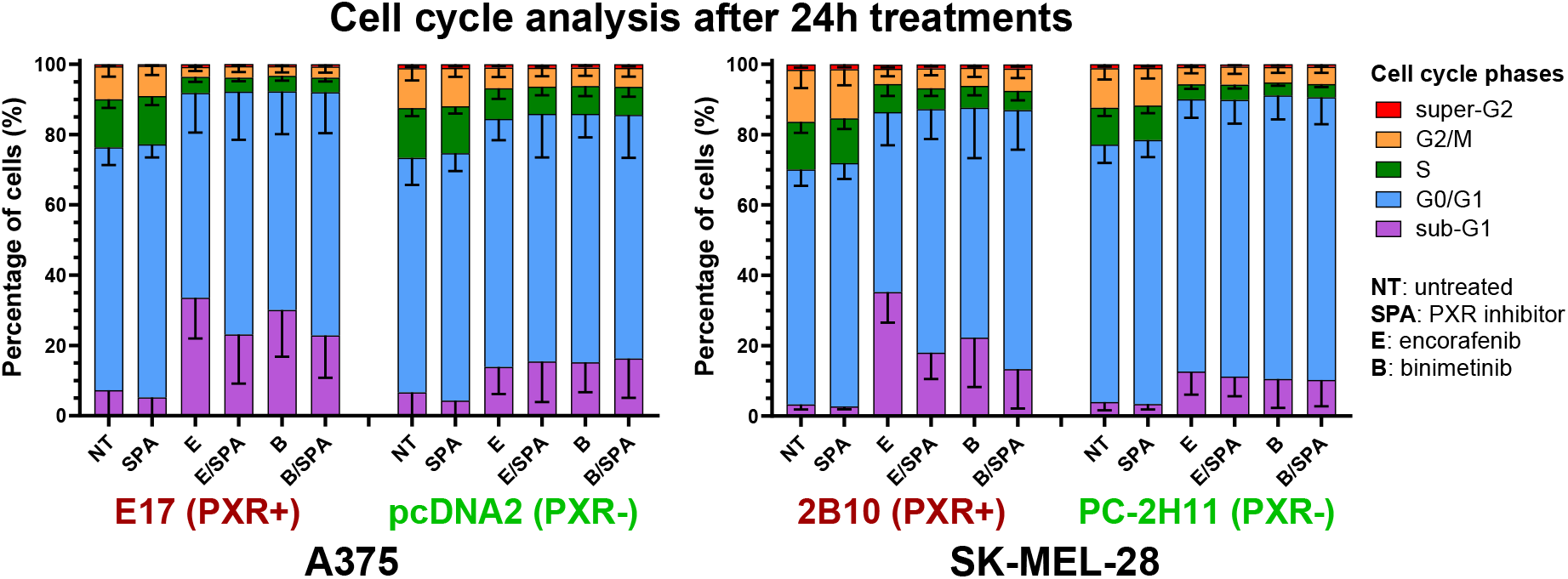
Cell cycle analysis following targeted therapy with encorafenib and binimetinib. DNA content measurement by propidium iodide staining after 24 h of treatment in PXR+ and PXR-clones derived from the A375 and SK-MEL-28 cell lines. Cells were treated with encorafenib (10 nM for A375, 40 nM for SK-MEL-28) and binimetinib (50 nM for A375, 100 nM for SK-MEL-28), either alone or in combination with the PXR antagonist SPA70 (1 μM).

**Figure S3.**
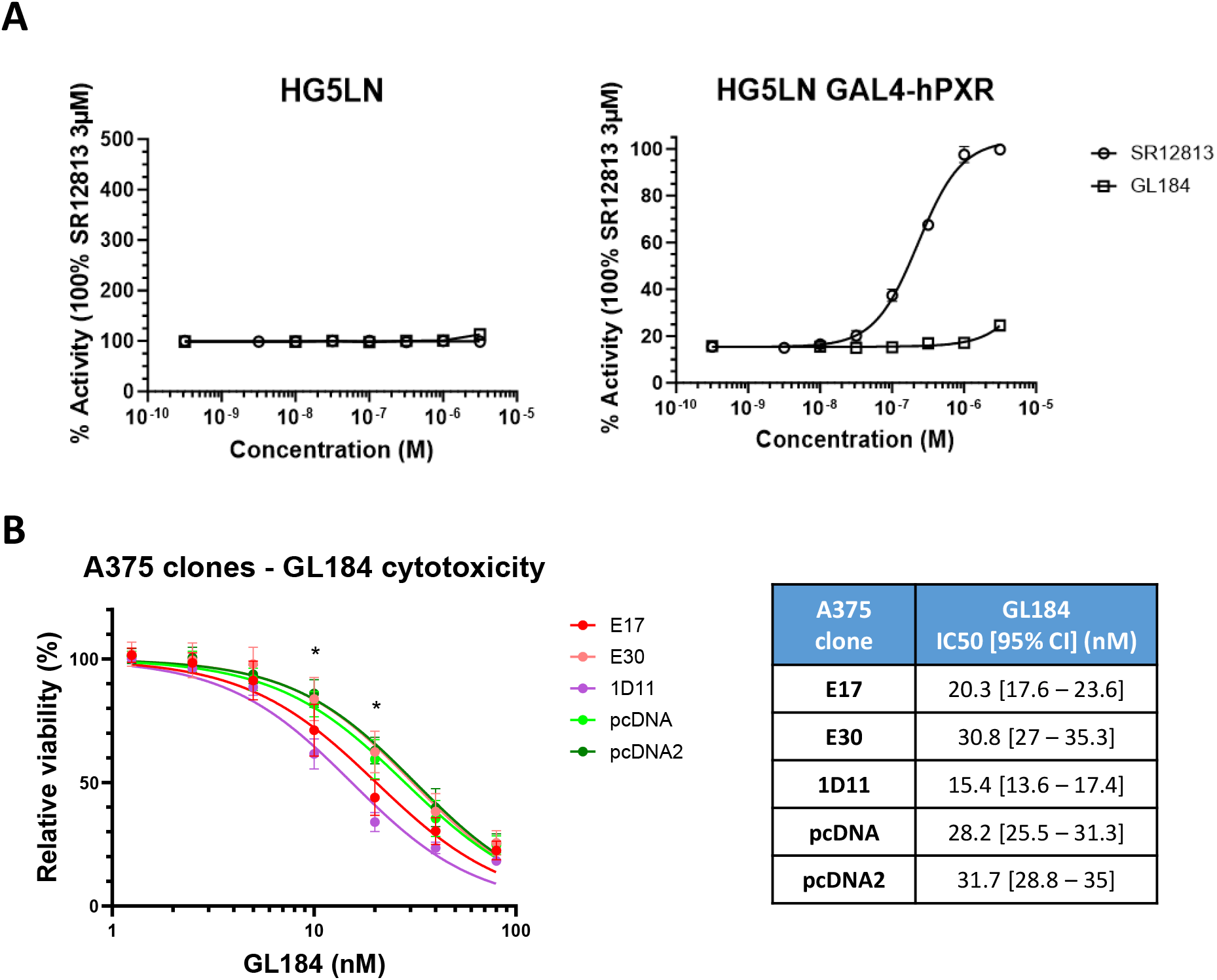
Analysis of GL184 activity on PXR and in A375 cells. **A)** Analysis of the agonistic activity of GL184 after 24 h of treatment using the HG5LN GAL4-hPXR luciferase reporter assay. **B)** Cytotoxic effects of GL184 in A375 clones, assessed using the sulforhodamine B assay after 72 h of treatment. Statistical significance was determined using the non-parametric Kruskal-Wallis test. ^∗^p value < 0.05, ^∗∗^p value < 0.01, ^∗∗∗^p value < 0.001.

**Figure S4.**
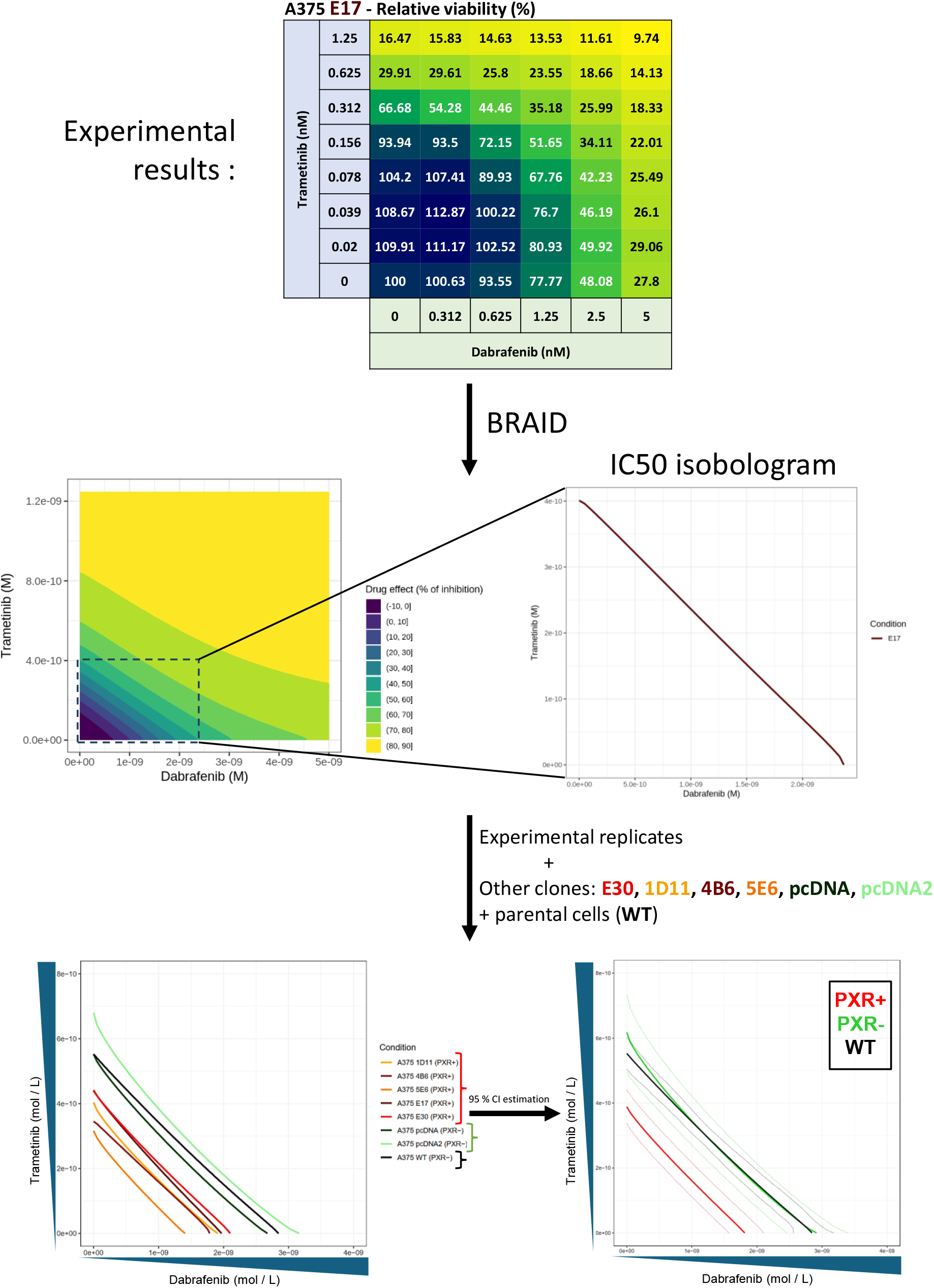
Workflow for the analysis of drug combination effects. BRAID (Bivariate Response to Additive Interacting Doses)^38^ was used to fit response surfaces to the experimental data and determine the drug concentrations required to achieve 50% growth inhibition (IC_50_) for each drug combination, which were subsequently used to generate isobolograms. Data from multiple clones and independent experimental replicates were combined to compare the fitted response surfaces and visualize differences in sensitivity to drug combinations. A representative replicate of the A375 E17 clone is shown as an example. For each cell group (PXR+, PXR−, or parental A375 WT), the mean response was calculated, and the corresponding 95% confidence interval was represented by dotted lines. The mean isobolograms obtained in the absence of SPA70 recapitulate the results shown in Fig. 4B.

**Figure S5.**
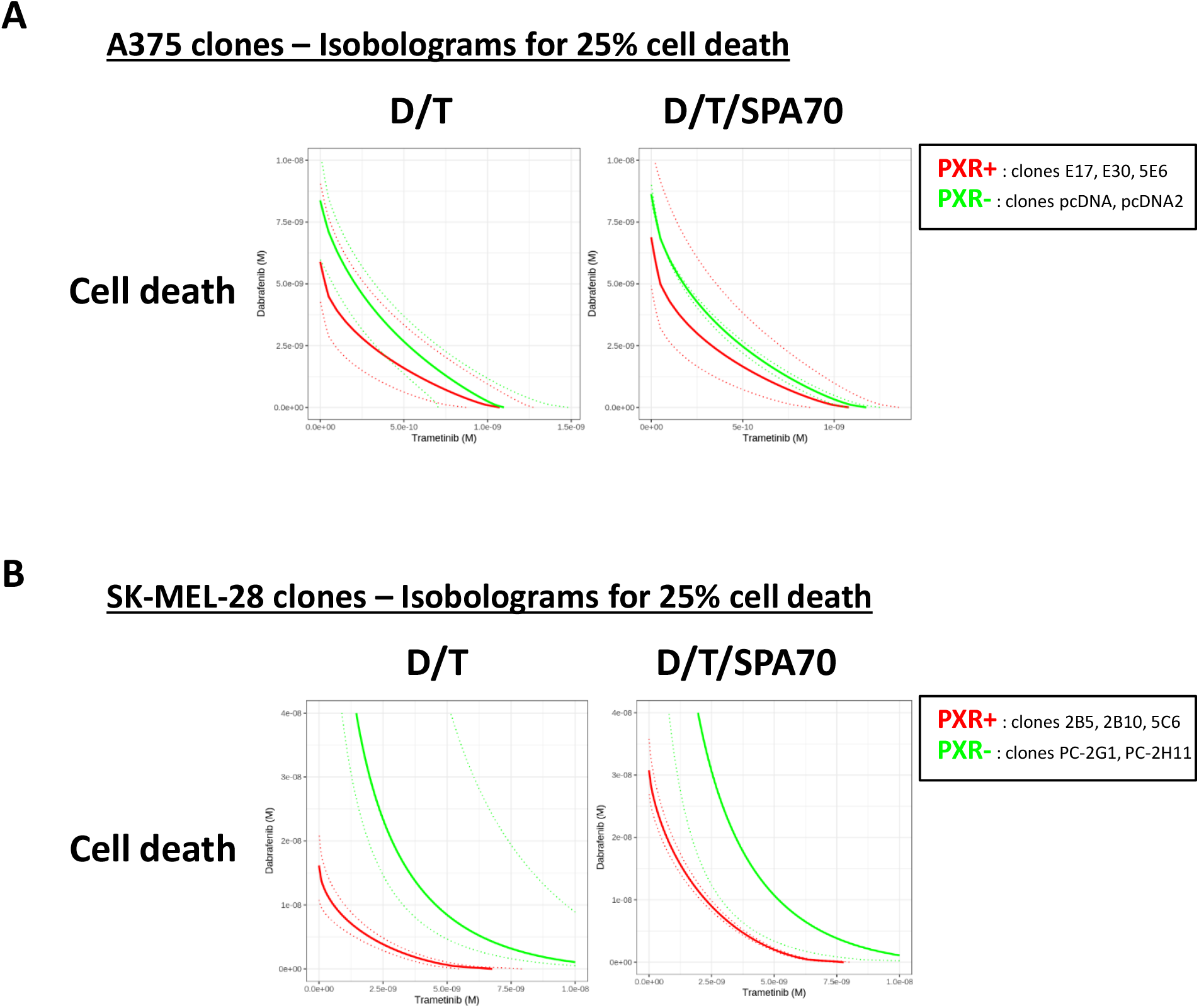
Effects of PXR expression and inhibition on melanoma cell death in response to combinations of BRAFi and MEKi in A375 and SK-MEL-28 cells. A) Isobolograms showing the effect of PXR overexpression in A375 clones on cell death following dabrafenib and trametinib treatment, in the presence or absence of the PXR antagonist SPA70 (1 μM), as determined by image analysis of propidium iodide-stained cells. Isobolograms represent drug combinations inducing 25% cell death. B) Same as in A) for SK-MEL-28 clones.

**Figure S6.**
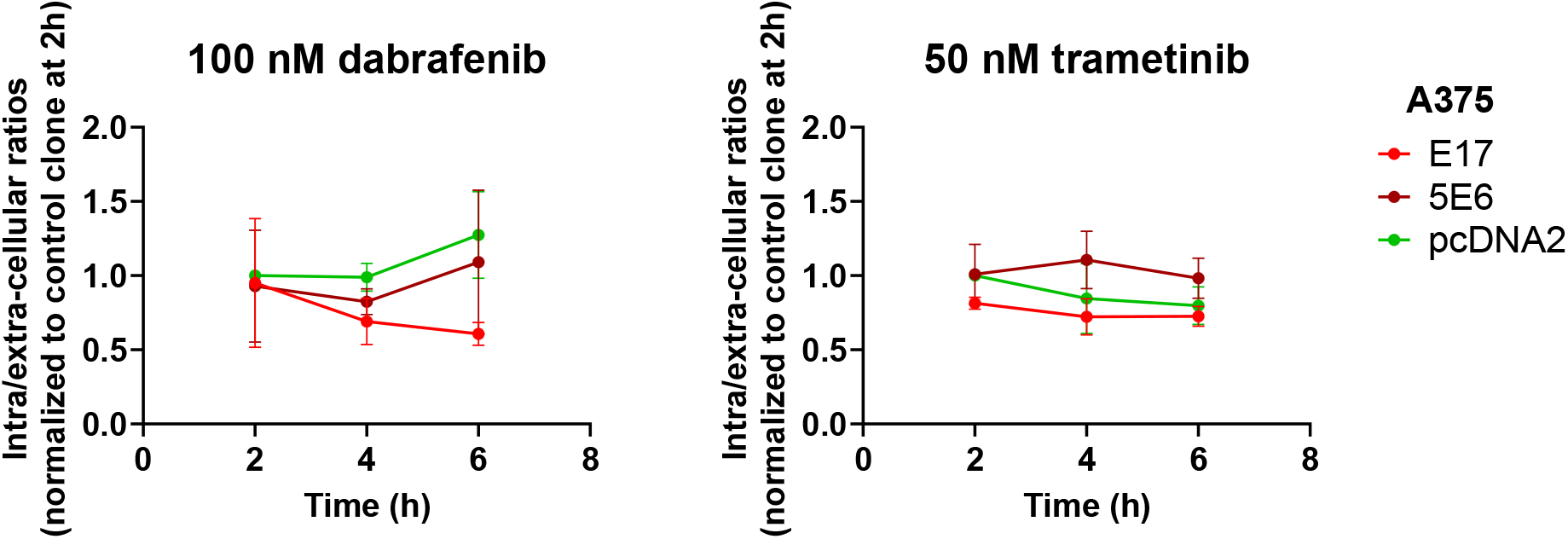
Analysis of dabrafenib and trametinib uptake by melanoma cells over 6. **h.** A375 clones stably expressing PXR (E17 and 5E6) and the control clone (pcDNA2) were treated with 100 nM dabrafenib or 50 nM trametinib for 2, 4, and 6 h. Drug concentrations were quantified by mass spectrometry, as described in the Materials and Methods section. Results are expressed as the intracellular/extracellular ratio of drug concentrations, normalized to the earliest time point of the control clone (pcDNA2), which was set to 1.

**Figure S7.**
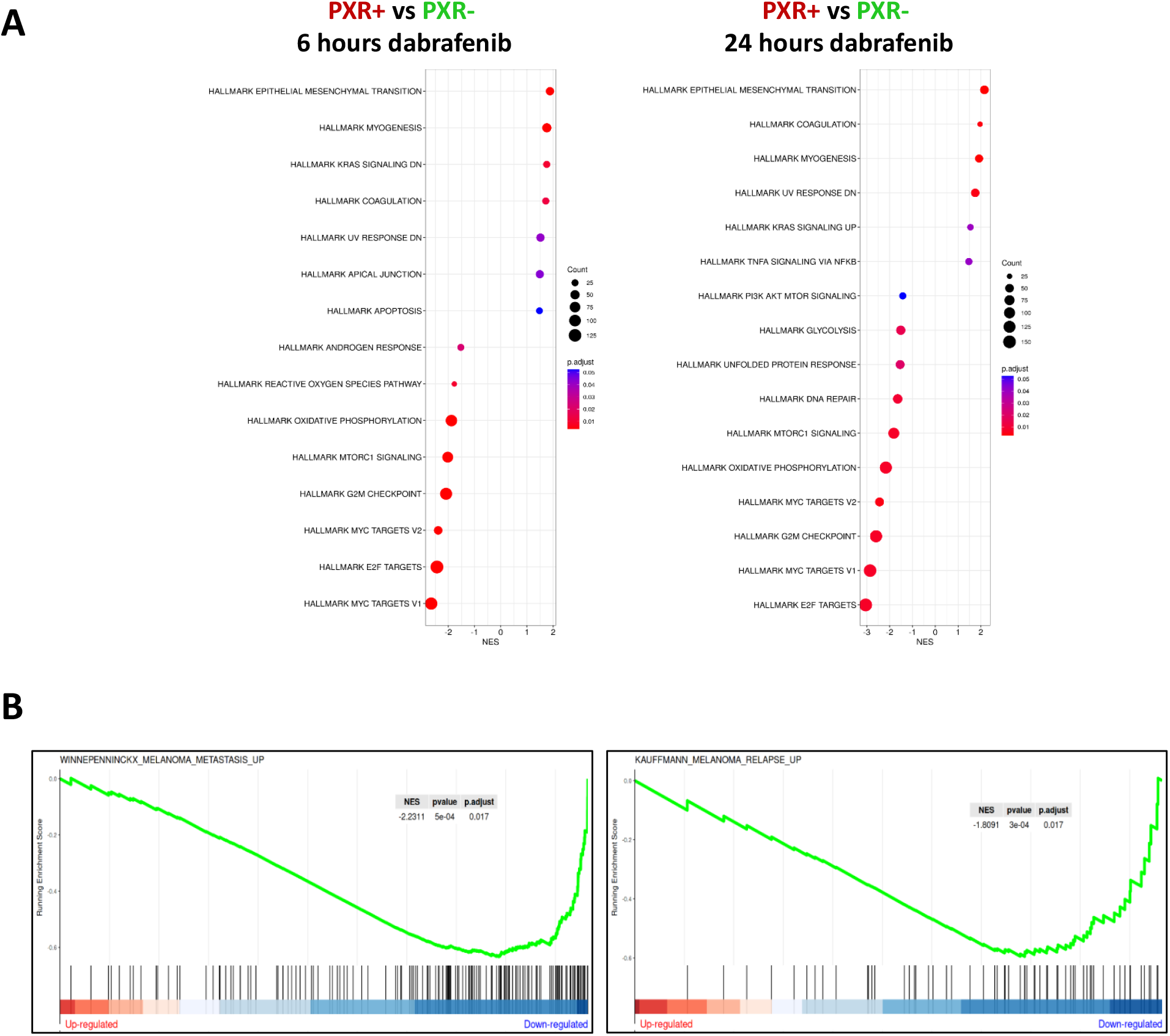
Gene Set Enrichment Analysis (GSEA) comparing A375 PXR-overexpressing clones with control clones. **A)** Gene Set Enrichment Analysis (GSEA) of Hallmark gene sets comparing A375 PXR+ clones with control clones following treatment with 10 nM dabrafenib for 6 or 24 h. Results are ranked according to the normalized enrichment score (NES). **B)** Gene Set Enrichment Analysis (GSEA) of gene sets defined by Winnepenninckx et al., 2006^42^ (WINNEPENNINCKX_MELANOMA_METASTASIS_UP) and Kauffmann et al., 2008^43^ (KAUFFMANN_MELANOMA_RELAPSE_UP), comparing A375 PXR-overexpressing clones with control clones under basal conditions. Both gene sets, whose upregulation has been associated with metastatic potential and melanoma relapse respectively, and thus with poor prognosis, were downregulated in PXR-overexpressing clones.

**Figure S8.**
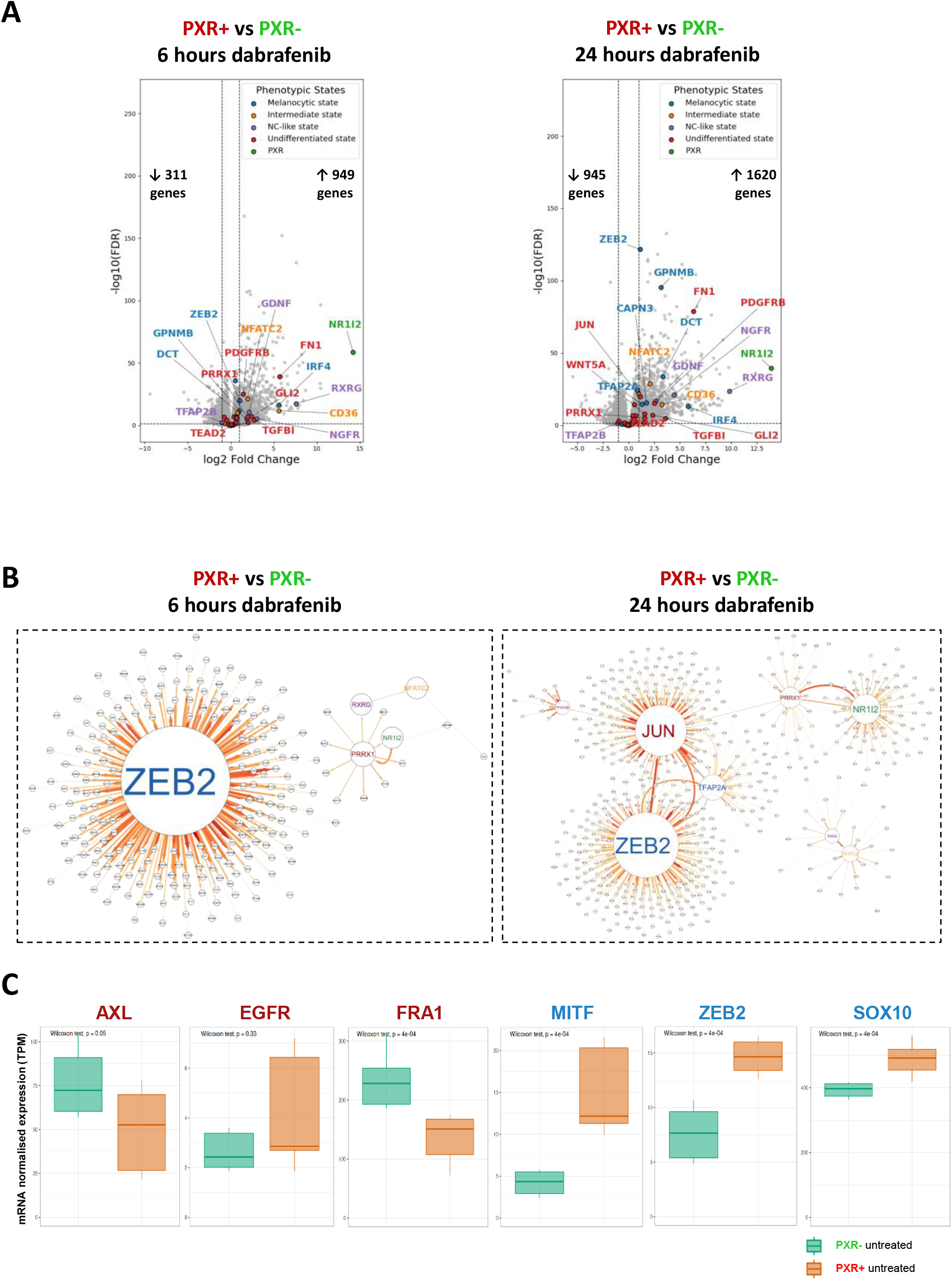
PXR-associated changes in melanoma phenotypic signatures. **A)** Volcano plots of differentially expressed melanoma phenotypic markers comparing A375 PXR+ clones with control clones following treatment with 10 nM dabrafenib for 6 or 24 h. Phenotypic markers, as defined by Pedri et al., 2022^13^, are color-coded according to their respective phenotypic state: melanocytic (blue), intermediate (orange), neural crest-like (NC-like, purple) and undifferentiated (red). NR1I2, the gene encoding PXR, is shown in green. Other genes are shown in gray. Numbers indicate the genes significantly upregulated (↑) or downregulated (↓) in PXR+ clones (|log2FC| ≥ 1 and FDR ≤ 0.05). **B)** Gene regulatory network inference analysis using GENIE3 under basal conditions or following treatment with 10 nM dabrafenib for 6 or 24 h. **C)** Normalized mRNA expression levels of AXL, EGFR, FRA1, MITF, ZEB2, and SOX10 under basal conditions in A375 clones. All markers showed |log2FC| < 1, with the exception of MITF. AXL and FRA1 were significantly downregulated, whereas MITF, ZEB2, and SOX10 were significantly upregulated in PXR-overexpressing clones. These markers were subsequently analyzed at the protein level by Western blotting. Statistical significance was determined using the non-parametric Wilcoxon test. ^∗^p value < 0.05, ^∗∗^p value < 0.01, ^∗∗∗^p value < 0.001.

**Figure S9.**
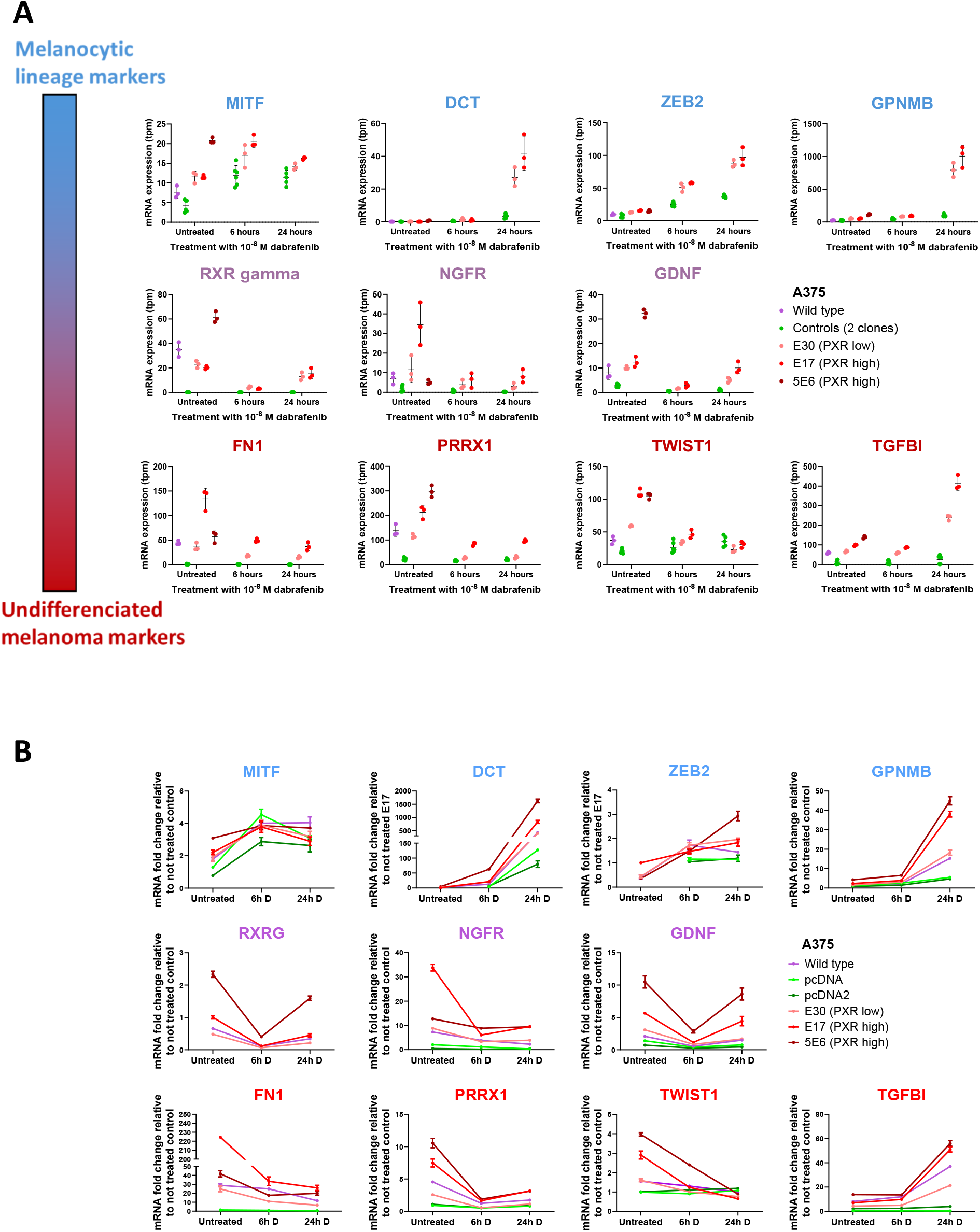
Analysis of phenotypic markers mRNA expression in A375 cells. **A)** Normalized expression (transcripts per million) of phenotypic markers in A375 cells, measured by RNA-seq, under basal conditions and following treatment with 10 nM dabrafenib for 6 or 24 h. **B)** RT-qPCR analysis of phenotypic marker expression in untreated A375 clones and following treatment with 10 nM dabrafenib for 6 h (**6h D**) or 24 h (**24h D**). Data are presented as mean fold change relative to the average expression of the corresponding untreated control clones ± SEM from one biological replicate. For DCT, ZEB2, and RXRG, which were expressed at levels too low to be quantified accurately in untreated control clones, fold change was calculated relative to untreated E17 clones ± SEM from one biological replicate.

**Figure S10.**
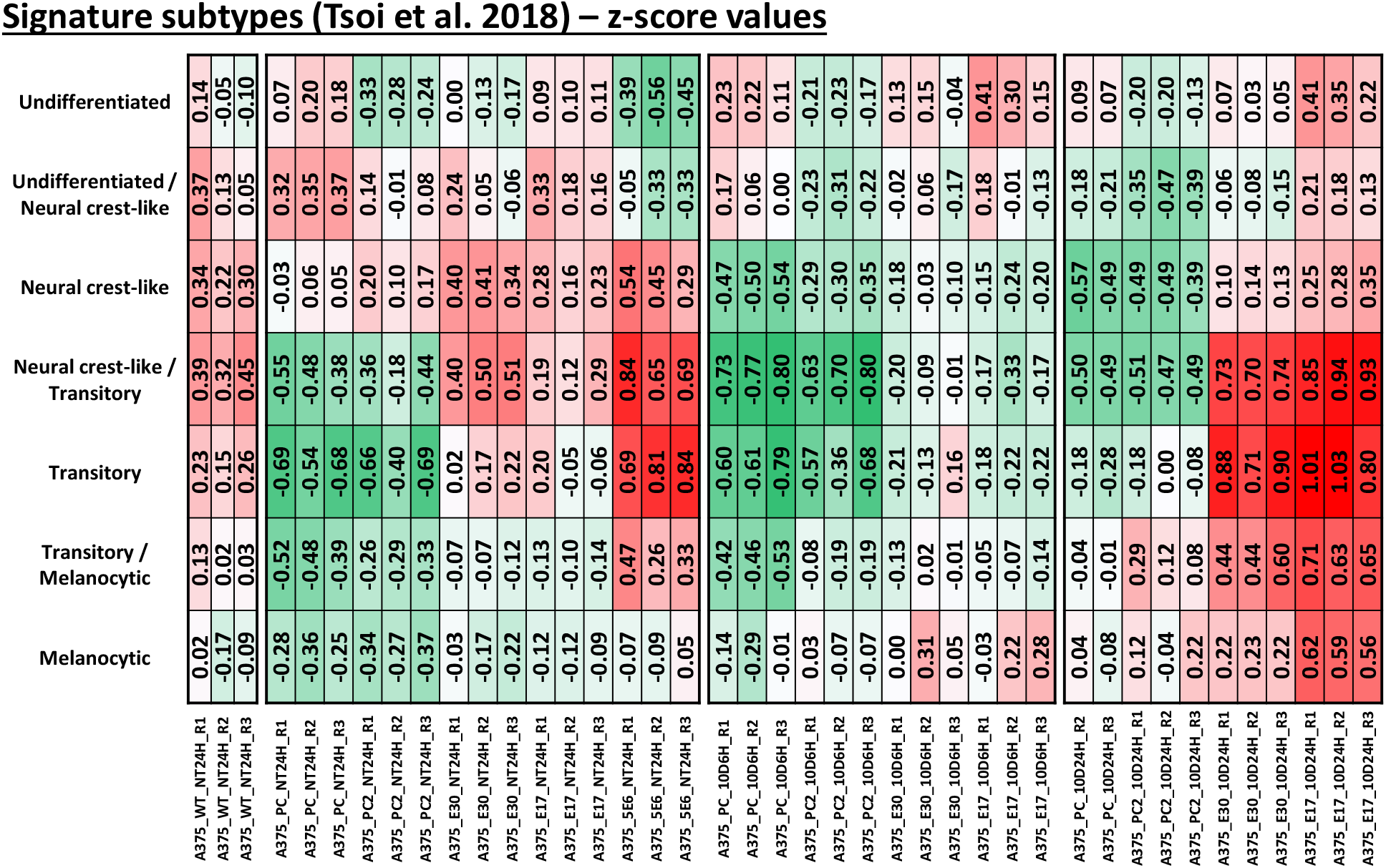
Numerical values of phenotypic signature z-scores for each sample of A375 clones. Sample organization and order are identical to those in Fig. 7B. Scaling was performed independently for each signature.

**Figure S11.**
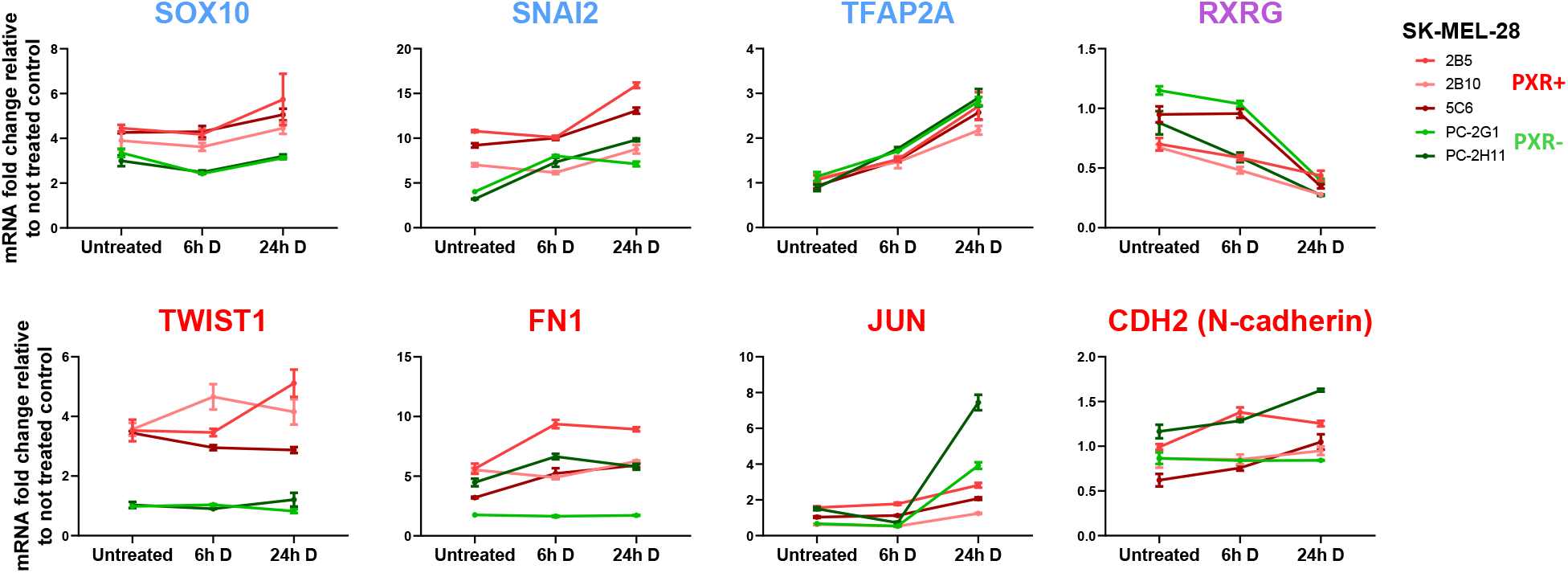
RT-qPCR analysis of phenotypic marker expression in untreated SK-MEL-28 clones and following treatment with 50 nM dabrafenib for 6 h (6h D) or 24 h (24h D). Data are presented as mean fold change relative to the average expression of untreated control clones ± SEM from one biological replicate.

**Figure S12.**
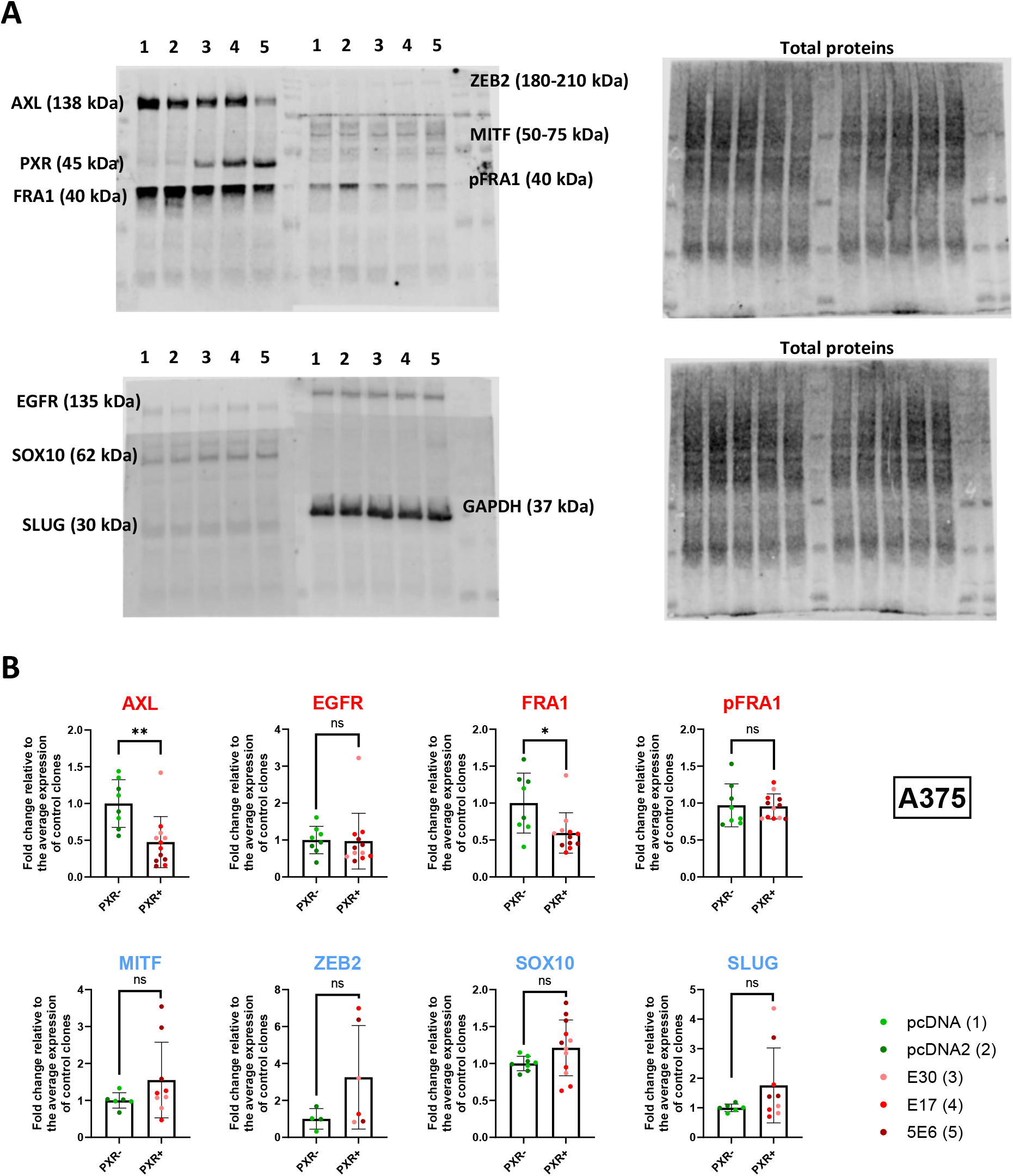
Western blot analysis of the expression of phenotypic markers in A375 clones. **A)** Uncropped Western blots showing the analysis of each phenotypic marker in A375 clones, with the corresponding molecular weights indicated. 1: pcDNA, 2: pcDNA2, 3: E30, 4: E17, 5: 5E6. Total protein staining was used as a loading control. **B)** Quantification of each phenotypic marker expression, normalized to the total protein signal and subsequently to the mean expression level of control clones, which was set to 1. At least two independent experimental replicates were performed using all five clones shown. Statistical significance was determined using the non-parametric Wilcoxon test. ^∗^p value < 0.05, ^∗∗^p value < 0.01, ^∗∗∗^p value < 0.001.

**Figure S13.**
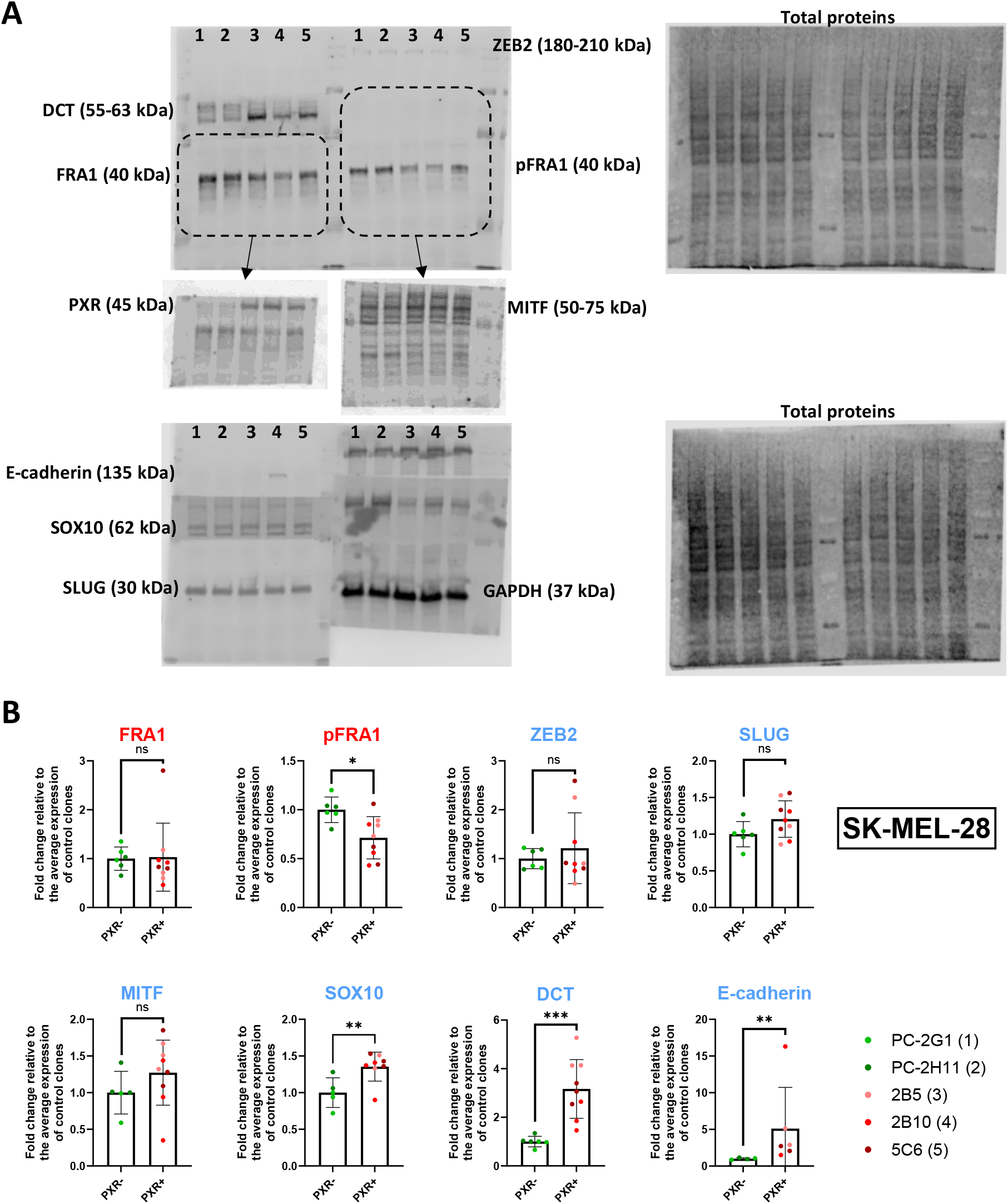
Western blot analysis of the expression of phenotypic markers in SK-MEL-28 clones. **A)** Uncropped Western blots showing the analysis of each phenotypic marker in SK-MEL-28 clones, with the corresponding molecular weights indicated. 1: PC-2G1, 2: PC-2H11, 3: 2B5, 4: 2B10, 5: 5C6. Total protein staining was used as a loading control. **B)** Quantification of each phenotypic marker expression, normalized to the total protein signal and subsequently to the mean expression level of control clones, which was set to 1. At least two independent experimental replicates were performed using all five clones shown. Statistical significance was determined using the non-parametric Wilcoxon test. ^∗^p value < 0.05, ^∗∗^p value < 0.01, ^∗∗∗^p value < 0.001.

---

**Supplementary Table 1.** Primary antibodies that were used for immunoblotting.

| **Antibody target** | **Clone** | **Host species** | **Catalog number** | **Supplier** | **Dilution** |
|---|---|---|---|---|---|
| PXR | G11 | Mouse | sc-48403 | Santa Cruz Biotechnology | 1:250 |
| GAPDH | ARC50888 | Rabbit | A19056 | ABclonal | 1:1000 |
| ABCB1 | E1Y7S | Rabbit | #13978 | Cell Signaling Technology | 1:1000 |
| ABCG2 | D5V2K | Rabbit | #42078 | Cell Signaling Technology | 1:1000 |
| MITF | D5G7V | Rabbit | #12590 | Cell Signaling Technology | 1:1000 |
| ZEB2 | E6U7Z | Rabbit | #97885 | Cell Signaling Technology | 1:1000 |
| DCT | — | Rabbit | ab74073 | Abcam | 1:1000 |
| SOX10 | EPR4007 | Rabbit | ab155279 | Abcam | 1:1000 |
| E-cadherin | 4A2 | Mouse | #14472 | Cell Signaling Technology | 1:1000 |
| SLUG | C19G7 | Rabbit | #9585 | Cell Signaling Technology | 1:1000 |
| EGFR | D38B1 | Rabbit | #4267 | Cell Signaling Technology | 1:1000 |
| AXL | C89E7 | Rabbit | #8661 | Cell Signaling Technology | 1:1000 |
| FRA1 | D80B4 | Rabbit | #5281 | Cell Signaling Technology | 1:1000 |
| Phospho-FRA1 (Ser265) | D22B1 | Rabbit | #5841 | Cell Signaling Technology | 1:1000 |

**Supplementary Table 2.**
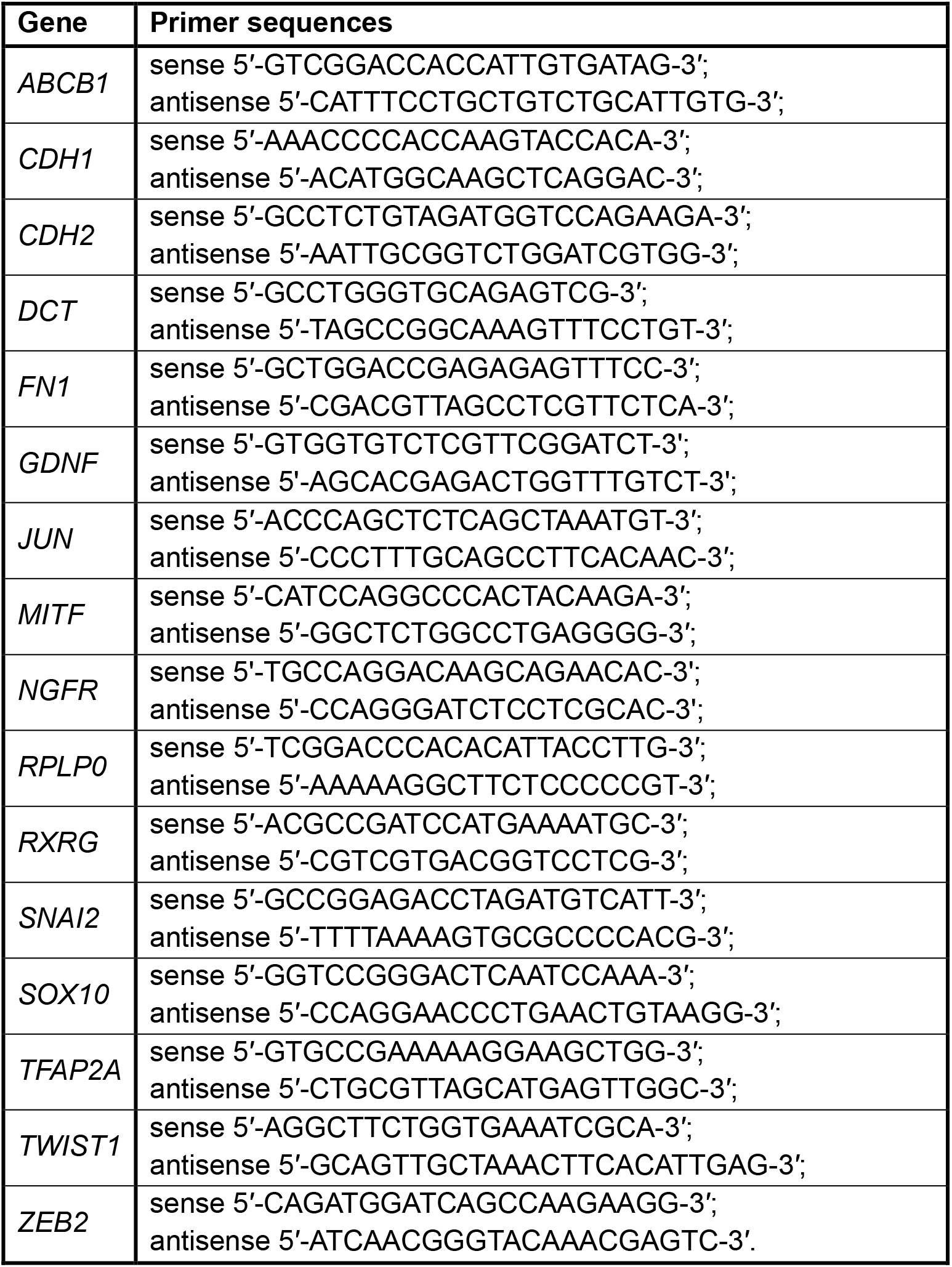
DNA primers that were used for RT-qPCR.

